# Single-molecule imaging reveals cytoplasmic translation of P granule-enriched mRNAs in *C. elegans*

**DOI:** 10.64898/2026.07.01.735846

**Authors:** William R Simmons, Qi Geng, Stephanie ITT Miller, Erik Griffin, Geraldine Seydoux

## Abstract

Germ granules are condensates in germ plasm, a specialized cytoplasm that segregates to the embryonic germline. In *Drosophila*, translation of *nanos* mRNA occurs at the surface of germ granules, suggesting that the granules promote translation. In *C. elegans*, however, germ (P) granules are not essential for Nanos expression. Using single-molecule imaging in *C. elegans* embryos, we map the distribution of translating and non-translating molecules of the Nanos homolog *nos-2* and two other maternal mRNAs enriched in P granules. In early germline blastomeres, these mRNAs are not translated and distribute between the cytoplasm and P granules. At translation onset, mRNA molecules in the cytoplasm are translated, while most mRNA molecules in the P granules remain non-translating. *nos-2* translation requires a rise in the concentration of the RNA-binding protein POS-1, which occurs independently of P granules. Consistent with low translation inside the granules, P granules are depleted of ribosomes and 43S pre-initiation complexes. Our observations suggest that germ granules promote Nanos protein expression by concentrating Nanos mRNA in germline precursors, but do not directly promote translation.

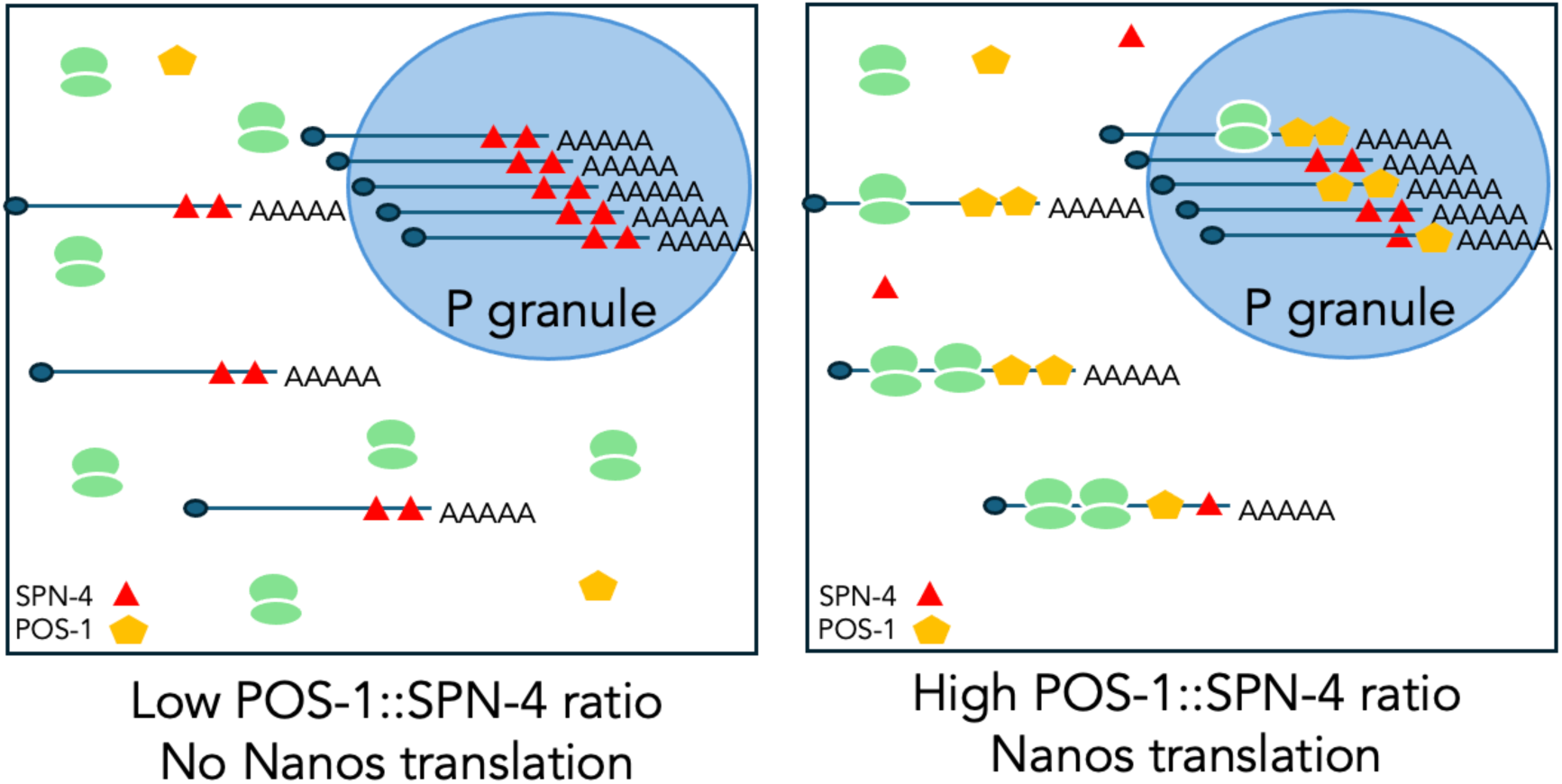

**Synopsis:** Germ granules are condensates proposed to regulate the translation of mRNAs like Nanos that code for germ cell fate determinants. Using single-molecule imaging in *C. elegans* embryos, this study shows that P granule scaffolds concentrate mRNAs in germline precursors, but do not control the activity of translational regulators.

- P granules concentrate mRNAs but are depleted of ribosomes and 43S pre-initiation complexes
- Translation occurs mainly in the cytoplasm where ribosomes are most abundant
*- nanos* translation onset is timed by a rise in POS-1, which counteracts the repressor SPN-4; both enrich in P granules but act independently.

## Introduction

RNA granules are condensates that concentrate mRNAs and have been proposed to regulate mRNA translation and/or stability in the cytoplasm (Ivanov *et al*, 2019; Chiappetta *et al*, 2022). For example, stress granules enrich translationally repressed mRNAs and arrested translation initiation complexes, suggestive of a function in translational repression (Kedersha *et al*, 1999). A challenge in determining the function of condensates, however, is that proteins enriched in condensates often also exist at a lower concentration in the cytoplasm, which complicates a determination of their primary site of action (Mateju & Chao, 2022; Putnam *et al*, 2023). Single-molecule techniques that visualize biological activity with high resolution have been used to address this challenge. Translation sites have been visualized using multi-copy epitope arrays (e.g. SunTag) that enable the *in situ* detection of nascent peptides emerging from ribosomes (Pichon *et al*, 2016; Wang *et al*, 2016; Wu *et al*, 2016; Yan *et al*, 2016). This method was used to demonstrate that certain mRNAs are translated inside stress granules, arguing against a simple repressive role for these condensates (Mateju *et al*, 2020).

Germ granules are another class of cytoplasmic condensates that have been implicated in translational control (Chiappetta *et al*, 2022). Germ granules assemble in germ plasm, a specialized cytoplasm that is segregated to the precursors of the germline during early embryogenesis (Hegner, 1908; Illmensee & Mahowald, 1974). Germ granules concentrate maternal proteins and mRNAs required for germ cell development, including *nanos* mRNA (Wang & Lehmann, 1991; Subramaniam & Seydoux, 1999; Köprunner *et al*, 2001; Juliano *et al*, 2010; MacArthur *et al*, 1999). Recently, SunTag analyses in *Drosophila* revealed that *nanos* mRNA molecules are often translated on the surface of polar granules (Chen *et al*, 2024; Ramat *et al*, 2024), raising the possibility that germ granules activate translation. Studies in *C. elegans*, in contrast, have suggested that germ granules are dispensable for *nanos* translation (Lee *et al*, 2020; Parker *et al*, 2020; Scholl *et al*, 2024; Cassani & Seydoux, 2022), but those studies did not determine where *nanos* translation occurs relative to granules.

In *C. elegans*, germ plasm assembles in the zygote and is asymmetrically segregated to the germ (P) lineage (Fig. S1A; Phillips & Updike, 2022). Proteins and mRNAs in germ plasm distribute between a soluble phase and two condensate types: P granules (analogous to the germ/polar granules of *Drosophila*) and germline P-bodies. P granules enrich Vasa family helicases, Argonautes and other small RNA machinery essential for gene regulation in primordial germ cells (PGCs) (Phillips & Updike, 2022). Germline P-bodies enrich mRNA regulators required for maternal mRNA turnover in PGCs (Cassani & Seydoux, 2024). P granules are stabilized in germ plasm by two paralogous intrinsically-disordered proteins MEG-3 and MEG-4 (Wang *et al*, 2014). MEG-3/4 recruit hundreds of maternal mRNAs to P granules, including the *nanos* homolog *nos-2* (Lee *et al*, 2020; Scholl *et al*, 2024). In *meg-3 meg-4* zygotes, P granules do not recruit mRNAs and dissolve by the 4-cell stage. The cytoplasmic pool of germ plasm proteins (and germline P-bodies), however, still assembles and *nos-2* mRNA is translated in the P_4_ founder cell (Wang *et al*, 2014; Lee *et al*, 2020; Cassani & Seydoux, 2022). NOS-2 protein levels, however, are lower in *meg-3 meg-4* embryos compared to wild-type (Cassani & Seydoux, 2022), raising the possibility that P granules could contribute to translational activation as reported in *Drosophila*.

In this study, we use single-molecule tools to examine this possibility by visualizing translation sites and quantifying mRNA and protein levels in wild-type and *meg-3 meg-4* mutants. We show that translation of *nos-2*, and two other P granule-enriched mRNAs, occurs in the cytoplasm, and only rarely in P granules which are depleted of ribosomes. MEG-3/4 increase the pool of mRNAs available for translation in P blastomeres but do not affect translational regulation, which depends on proteins that enrich in germ plasm independently of MEG-3/4. Our findings potentially reconcile observations in *Drosophila* and *C. elegans* by suggesting that germ granule scaffolds boost Nanos expression by increasing local mRNA concentration, but do not regulate translation, which depends on factors that enrich in the granules but function independently.

## Results

### Tagging strategy

Our strategy to visualize translation sites is modeled after existing methods (Blake *et al*, 2024; van der Salm *et al*, 2025) that label nascent peptides using tandem arrays of peptide epitopes inserted at the N-terminus of the protein of interest coupled with smFISH to detect transcripts (Fig. S1B). Using CRISPR genome editing (Paix *et al*, 2017), we inserted a multi-copy tag (7xHA or 8xALFA-tag) in three loci known to produce maternal transcripts that enrich in P granules (*nos-2*, *xnd-1* and *zif-1*; Scholl et al, 2024) (Materials and Methods).

Prior reports described the expression pattern of NOS-2, XND-1 and ZIF-1 proteins using 3’ UTR reporters and/or antibodies (Fig. S1C). NOS-2 and XND-1 are germline-specific proteins that accumulate in the germline founder cell P_4_ and its immediate daughters Z2 and Z3. NOS-2 localizes in the cytoplasm and on P granules (Subramaniam & Seydoux, 1999; Lee *et al*, 2020), whereas XND-1 is a nuclear protein (Mainpal *et al*, 2015). ZIF-1 is an E3 ubiquitin ligase complex adaptor predicted to be expressed in the cytoplasm of somatic blastomeres and of the primordial germ cells Z2 and Z3 (DeRenzo *et al*, 2003; Schwartz *et al*, 2023; Guven-Ozkan *et al*, 2010). ZIF-1 has a short half-life and a GFP::ZIF-1 fusion could only be detected when other subunits of the E3 complex were depleted by RNAi (DeRenzo *et al*, 2003). Immunofluorescence detection using anti-HA and anti-ALFA antibodies in fixed embryos confirmed that the tagged proteins were expressed with the correct lineage specificity and subcellular localization (Fig. EV1).

### *meg-3 meg-4* mutants lacking P granules initiate translation of *nos-2*, *xnd-1* and *zif-1* mRNAs with wild-type timing

*meg-3 meg-4* mutants accumulate lower levels of NOS-2 and XND-1 proteins in P_4_ blastomeres (Cassani & Seydoux, 2022), which could be due to delayed translation onset, lower mRNA levels, and/or lower translation efficiency. To determine the timing of translation onset in the P lineage, we immunostained 7xHA::NOS-2, 8xALFA::XND-1 and 7xHA::ZIF-1 in wild-type embryos and embryos derived from *meg-3(ax3055) meg-4(ax3052)* mutant hermaphrodites (hereafter referred to as *meg-3 meg-4* embryos) and determined when the signal in germline blastomeres exceeds background levels (Materials and Methods). We first detected 7xHA::NOS-2 and 8xALFA::XND-1 in P_3_ in 12-to-15-cell stage embryos and 7xHA::ZIF-1 in P_4_ in ∼50-cell stage embryos (Fig. 1A). These stages are one cell cycle earlier than prior estimates reported in the literature, consistent with the higher sensitivity conferred by the multi-copy tags (Fig. S1C). We observed the same timing of translational onset in *meg-3 meg-4* embryos (Fig. 1A). Quantification of 7xHA::NOS-2 levels at different stages confirmed that NOS-2 protein accumulation initiates in P_3_ in *meg-3 meg-4* embryos as in wild-type (Fig. 1B). The only difference is that less NOS-2 protein accumulates over the lifetime of P_4_.

**Figure 1:**
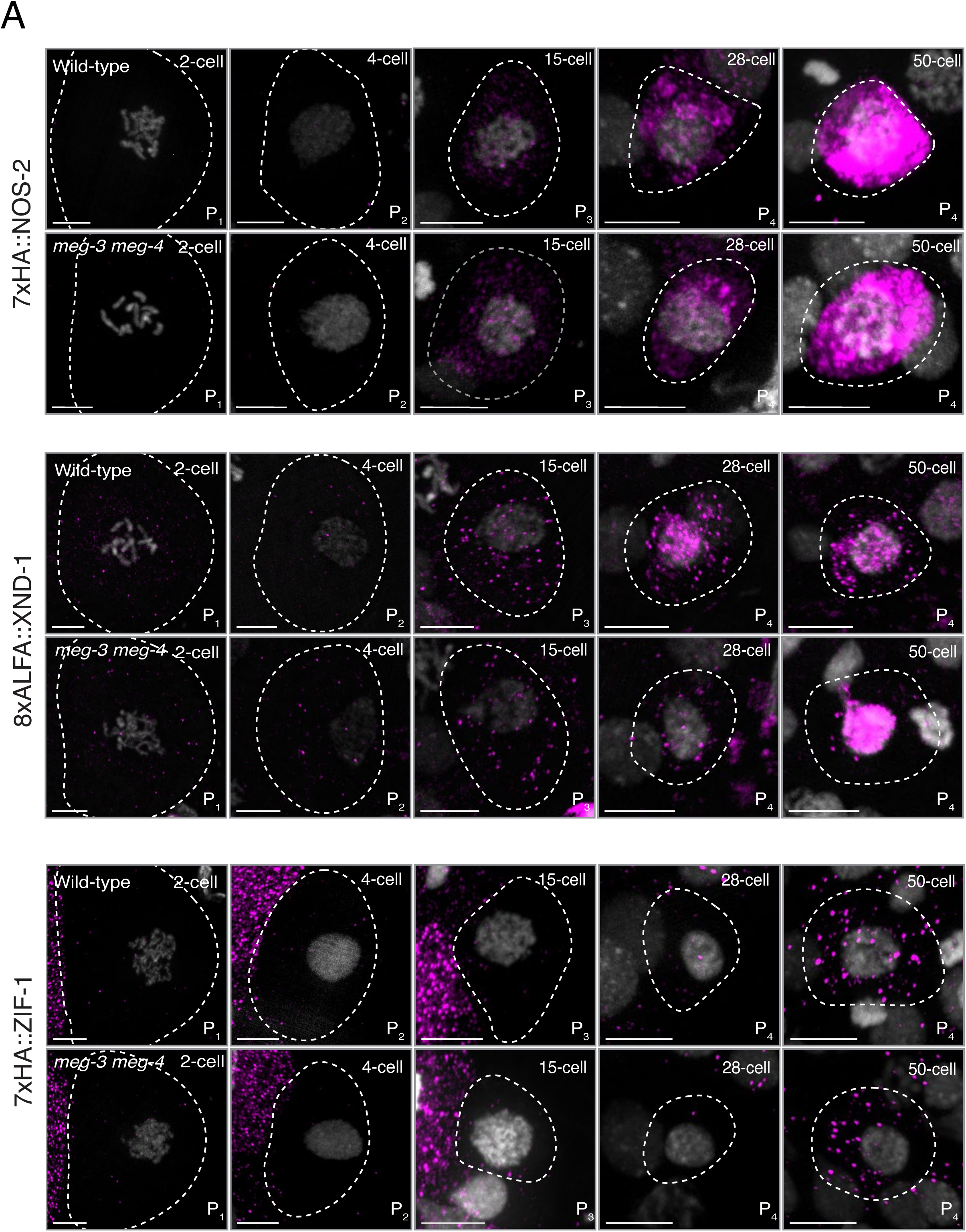
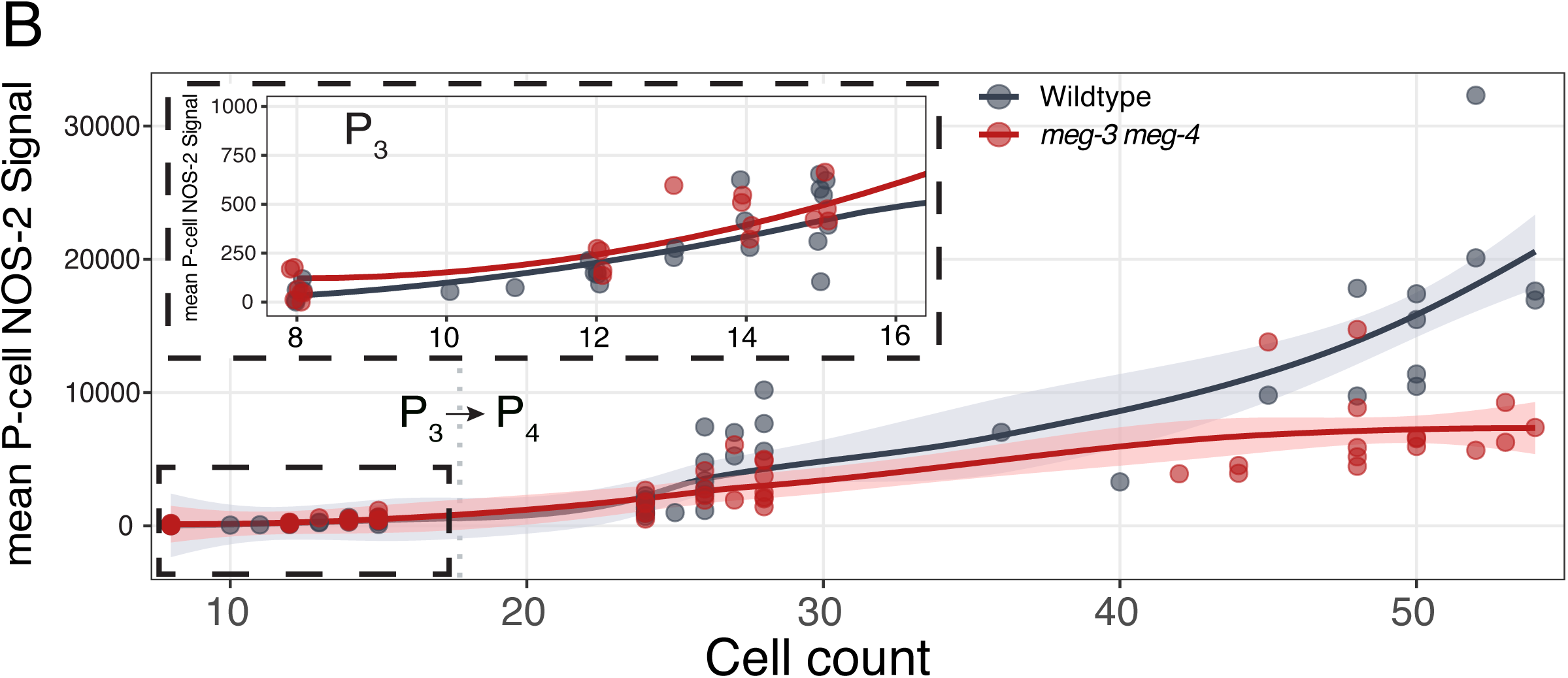
Translation onset of NOS-2, XND-1 and ZIF-1 occurs with the same timing in *w*ild-type and *meg-3 meg-4* mutants. **(A)** Photomicrographs of P blastomeres immunostained for the fusion proteins indicated comparing wild-type and *meg-3(ax3055) meg-4(ax3052)* mutants. Dashed lines represent cell boundaries. Images are scaled to accommodate the diminishing sizes of P_1_ through P_4_. Scale bars are 5µm. **(B)** Graph showing 7xHA::NOS-2 signal intensity (arbitrary units, Y axis) in P blastomeres normalized to background signal in somatic blastomeres (0) comparing wild-type and *meg-3(ax3055) meg-4(ax3052)* mutants at different embryonic stages (X-axis). The inset expands early stages. Each point is a single P blastomere. Lines are loess smoothed curve of the mean. Shaded areas represent 95% confidence interval.

### Cytoplasmic levels of *nos-2* and *xnd-1* mRNAs in P_4_ are lower in *meg-3 meg-4* mutants compared to wild-type

Prior studies reported that, on average, the concentration of *nos-*2 and *xnd-1* mRNAs is reduced in P_4_ in *meg-3 meg-4* embryos compared to wild-type, but these studies did not distinguish between mRNAs in the cytoplasm and in granules. To address this, we developed a single-molecule spot detection pipeline to count individual transcripts in the cytoplasm and in granules as detected by single-molecule fluorescence *in situ* hybridization (smFISH) in Z-stacks spanning entire P_3_ and P_4_ blastomeres (Fig. EV2). *meg-3 meg-4* embryos lack P granules but maintain germline P-bodies, which are smaller condensates (Cassani & Seydoux, 2022). We used a probe against the SL1 splice leader sequence present on the 5’ end of approximately 70% of *C. elegans* mRNAs (Allen *et al*, 2011) to mark both P granules and germline P-bodies (Cassani & Seydoux, 2022). Consistent with the lack of P granules, overall mRNA counts of *nos-2*, *xnd-1* and *zif-1* were lower in P_4_ blastomeres in *meg-3 meg-4* embryos compared to wild-type (Fig. 2). P_4_ blastomeres are approximately 1/3 of the volume of P_3_ blastomeres and therefore should contain 1/3 of the mRNA molecules observed in P_3_ if mRNAs are partitioned in an unbiased manner at the P_3_ division. We observed the expected levels in P_4_ relative to P_3_ in *meg-3 meg-4* embryos but higher levels in wild-type embryos (Fig. 2), consistent with asymmetric segregation of P granules at the P_3_ division (Fig. S1A). Interestingly, in wild-type embryos, the cytoplasmic levels of *nos-2* and *xnd-1* (but not *zif-1*) were also higher in P_4_ than expected, indicating that MEG-3/4 increases both the cytoplasmic and granular pools of these transcripts in P_4_ (Fig. 2). The cytoplasmic pool was maintained during the lifetime of P_4_, but the granule-associated pool decreased by the 50-cell stage (Fig. 2). We conclude that 1. P_4_ blastomeres inherit lower levels of cytoplasmic *nos-2* and *xnd-1* mRNAs in *meg-3 meg-4* mutants compared to wild-type and 2. the pool of mRNAs associated with granules in wild-type embryos diminishes over the lifetime of P_4_.

**Figure 2:**
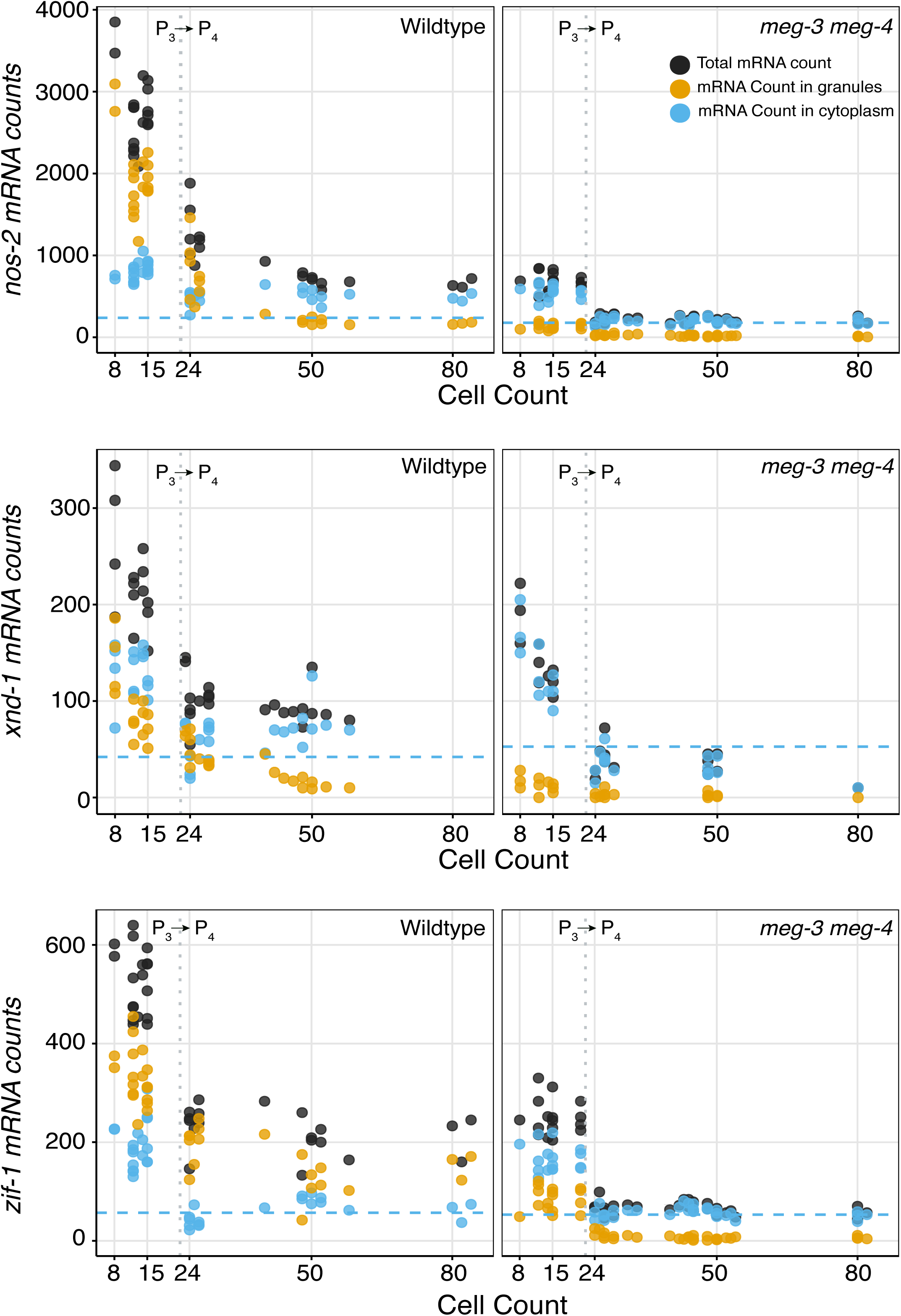
MEG-3 and MEG-4 bias mRNA segregation to P blastomeres. **(A)** Graphs showing the number of mRNA molecules (Y axis) in P_3_ and P_4_ blastomeres in different stages (X axis) comparing wild-type and *meg-3(ax3055) meg-4(ax3052)* embryos. Each point represents total (black), granular (gold) or cytoplasmic (blue) mRNA counts for one P blastomere. The blue stippled line shows the mRNA count expected in the cytoplasm if mRNAs segregate with no spatial bias at the P_3_ to P_4_ division. *nos-2* and *zif-1* counts were obtained from wild-type untagged animals, and *xnd-1* counts were obtained in animals expressing *8xALFA::xnd-1*.

### Translation occurs primarily in the cytoplasm of P blastomeres and does not require P granules

To determine the number and position of translating mRNAs in P blastomeres, we used our single-molecule spot detection pipeline to detect mRNAs colocalized with protein spots in images combining smFISH detection of mRNA with immunofluorescence detection of protein (Materials and Methods, Fig. 3). To verify that co-localized protein and RNA spots represent translation sites, we blocked translation using a 30min heat-shock (Fig. EV3). In embryos expressing 8xALFA ::XND-1, 26.1% (152 of 582) of *xnd-1* transcripts co-localized with protein spots under control conditions but only 0.37% (2 of 537) under heat shock conditions (Fig. EV3). Importantly, mature XND-1 protein could still be detected in nuclei under heat shock conditions, indicating that the treatment specifically eliminated nascent proteins (Fig. EV3). We conclude that mRNA spots that colocalize with protein spots correspond to translating mRNA molecules.

**Figure 3:**
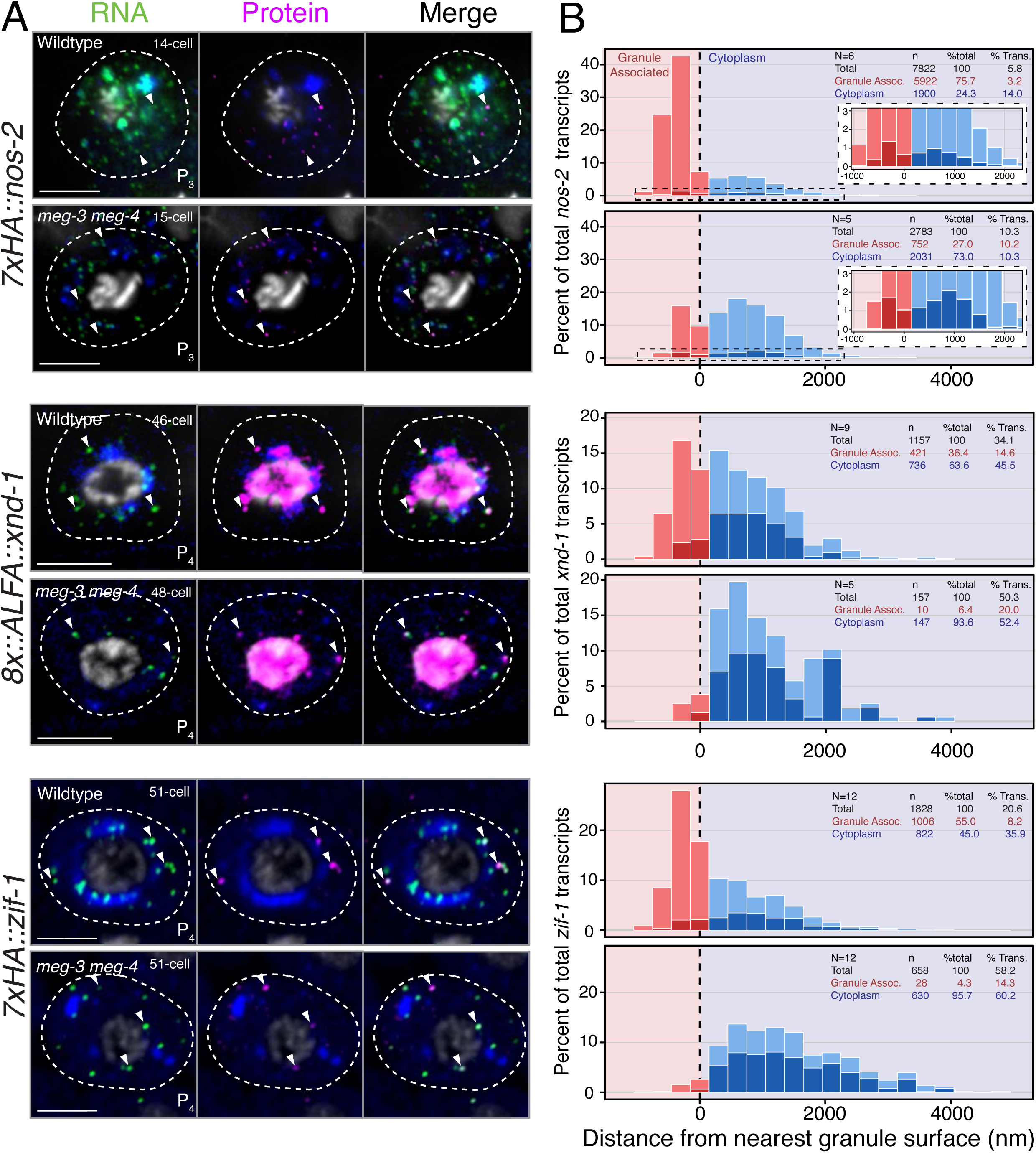
Most translating mRNAs are in the cytoplasm and do not require P granules. **(A)** Photomicrographs (single Z slices) of representative P_3_ blastomeres (*nos-2*) or P_4_ blastomeres (*zif-1* and *xnd-1*) in wild-type and *meg-3(ax3055) meg-4(ax3052)* embryos processed for immunostaining and smFISH (Materials and Methods). Green: RNA, Protein: Magenta, Blue: Granules (SL1), White: DNA. Dashed lines are cell boundaries. Arrow heads point to representative RNA spots co-localized with protein (translation spots). Scale bars are 5µm. **(B)** Stacked histograms showing the distribution of mRNAs relative to the surface of the nearest germ granule. mRNAs were counted in 300nm bins (x = 0, dashed vertical line). Negative values represent mRNAs inside granules (red), and positive values represent mRNAs outside granules (blue). Dark shading indicates translating mRNAs. Statistics show N = number of P blastomeres scored, n = number of mRNA molecules scored, % Trans = Percent translating mRNAs in each compartment. See Fig. EV3C for a different representation of these data showing mRNA counts for each P blastomere examined.

Because NOS-2 protein accumulates on P granules (Lee *et al*, 2020), we quantified *nos-2* translation sites at translation onset in P_3_ to minimize interference from mature protein. To define the pool of RNA in germ granules, we segmented granules using the SL1 signal, and accounting for the resolution limits of our imaging system, classified any mRNA spot within 150 nm of the SL1 boundary as granule associated. At this stage, 75.7% of all *nos-2* transcripts were granule associated (Fig. 3, Fig. EV3C). We found that only 3.2% of these transcripts were translating. In contrast, 14.0% of *nos-2* transcripts in the cytoplasm were translating. Translating mRNAs were distributed throughout the cytoplasm with no bias toward the granules (Fig. 3). As described above, *meg-3 meg-4* embryos have fewer mRNA molecules in the cytoplasm of P blastomeres compared to wild-type, with only a minority localizing to germline P-bodies.

Remarkably, despite the lower mRNA levels, the percent of translating transcripts in *meg-3 meg-4* mutants (10.3%) was similar to that observed for the cytoplasmic pool in wild-type (14.0%).

Unlike NOS-2, mature XND-1 and ZIF-1 proteins do not accumulate in the cytoplasm or on P granules, which allowed us to quantify translation sites when these proteins are robustly translated in P_4_ blastomeres (Fig. 3). Again, we observed a strong bias for translating transcripts to be in the cytoplasm (Fig. 3). For example, 35.9% of *zif-1* transcripts in the cytoplasm were translating compared to 8.2% associated with germ granules. The distribution of translating transcripts in the cytoplasm was not biased with respect to granules. Translation rates in *meg-3 meg-4* were similar (*xnd-1*) and even higher (*zif-1*) than that observed for the cytoplasmic pool in wild-type (Fig. 3).

We conclude that the majority of translating *nos-2*, *xnd-1* and *zif-1* molecules are in the cytoplasm. Cytoplasmic transcripts are translated with similar efficiencies in the presence or absence of P granules. Transcripts associated with germ granules are rarely translated.

### Translational regulation of *nos-2* correlates with increasing levels of POS-1 in the P lineage independent of P granules

*nos-2* translation is regulated by cis-acting elements in its 3’ UTR which contain binding sites for the repressor SPN-4 and the translational de-repressor POS-1 (Jadhav et al, 2008). Jadhav *et al*, 2008 reported that POS-1 competes with SPN-4 for binding to the *nos-2* 3’ UTR *in vitro* and suggested that POS-1 de-represses *nos-2* translation when its concentration overcomes that of SPN-4 in P_4_. To examine this possibility, we quantified the concentration of POS-1 and SPN-4 at different embryonic stages using GFP-tagged alleles. We found that POS-1 concentration increases over time in the P lineage, in contrast to SPN-4 which remains constant (Fig. 4). The ratio of POS-1 to SPN-4 concentration reaches 1.52 in P_3_, when *nos-2* translation is first activated (Fig. 1).

**Fig. 4:**
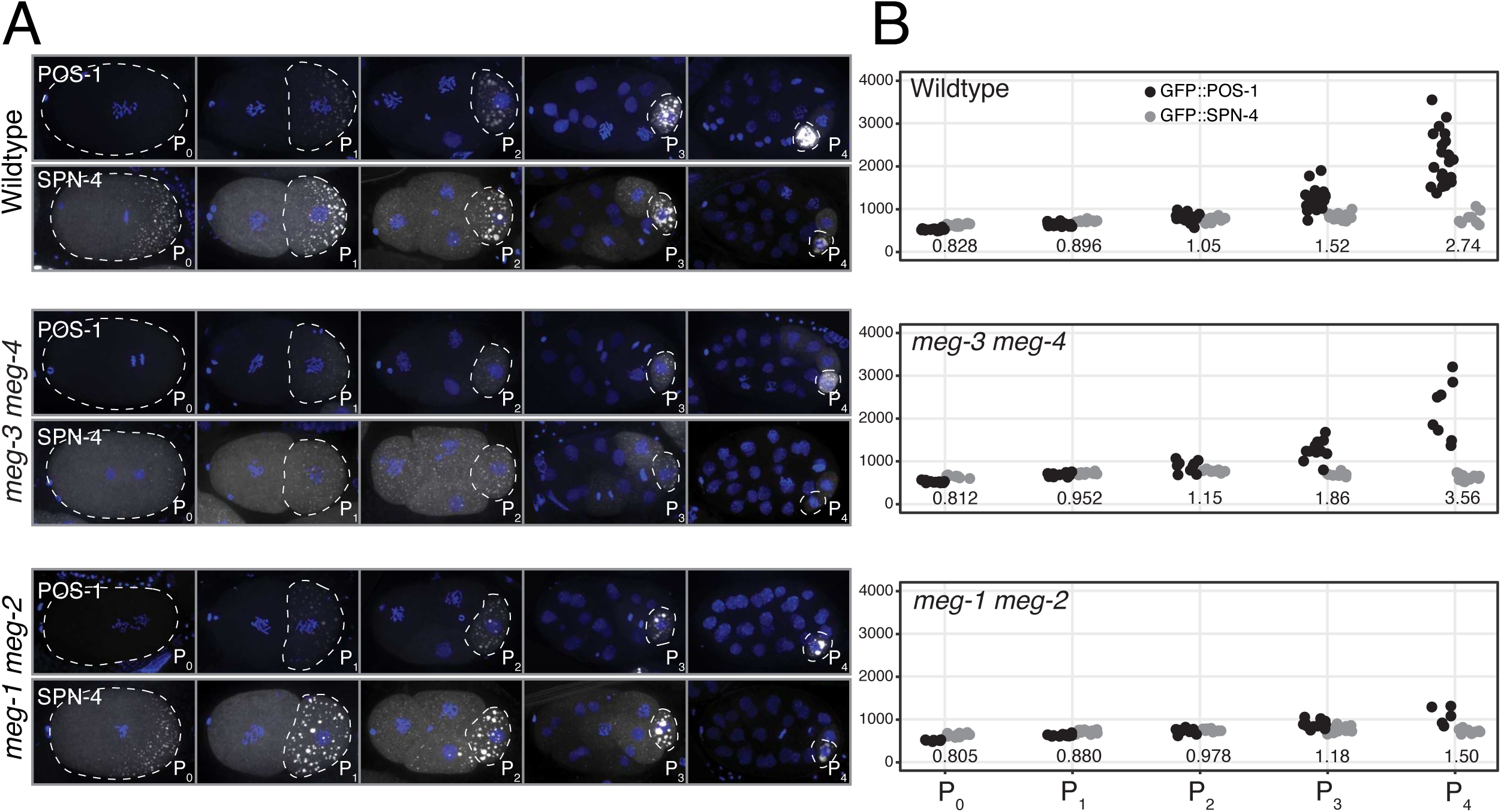
POS-1 levels increase in the P lineage independent of P granules. **(A)** Photomicrographs of embryos expressing either GFP::POS-1 or SPN-4::GFP in wild-type and germline-granule-deficient backgrounds. GFP::POS-1 was imaged in wild-type, *meg-3(ax3055) meg-4(ax3052)* mutant, and *meg-1/2(RNAi)* embryos. SPN-4::GFP was imaged in wild-type, *meg-3/4(RNAi)*, and *meg-1/2(RNAi)* embryos. Dashed lines outline each P blastomere. (B) Graphs showing the mean GFP signal (arbitrary units on Y axis) in P blastomeres. Each dot represents a single P blastomere scored for GFP::POS-1 (black) or SPN-4::GFP (grey). Numbers indicate the mean POS-1:SPN-4 ratio for each stage.

POS-1 and SPN-4 are germ plasm proteins that distribute between P granules, germline P-bodies and the cytoplasm. To explore what determines the relative concentrations of POS-1 and SPN-4, we examined POS-1 and SPN-4 levels in *meg-3 meg-4* and *meg-1 meg-2* mutants. MEG-1 (and its paralog MEG-2) are germ plasm proteins that co-immunoprecipitate with POS-1 and SPN-4 (Cassani & Seydoux, 2022). *meg-1 meg-2* mutants assemble normal P granules, but accumulate reduced levels of NOS-2 protein (Leacock & Reinke, 2008; Cassani & Seydoux, 2022). We found that SPN-4 levels were unaffected in *meg-3 meg-4* and *meg-1 meg-2* embryos. POS-1 levels in the P lineage increased in *meg-3 meg-4* embryos as in wild-type (Fig. 4). POS-1 levels were similar to wild-type in *meg-1 meg-2* zygotes but increased more slowly, reaching a POS-1:SPN-4 ratio of 1.50 only in P_4_ (Fig. 4). Interestingly, this is also the stage where we first detected 7xHA::NOS-2 accumulation in *meg-1 meg-2* embryos (Fig. EV4A and B), suggesting that the delayed POS-1 accumulation leads to a delay in *nos-2* translation onset. Knocking down *spn-4* by RNA-mediated interference in *meg-1 meg-2* embryos rescued 7xHA::NOS-2 accumulation in P_3_ blastomeres (Fig. EV4C and D). We conclude that, as proposed by Jadhav et al. (2008), *nos-2* translation onset requires an increase in POS-1 levels relative to SPN-4. This increase occurs independently of P granules and requires the germ plasm factors MEG-1 and MEG-2.

### Ribosomes and associated translation initiation factors are depleted from P granules

The observation that RNAs in granules are rarely translated raises the possibility that factors required for translation might be depleted from P granules. To examine this possibility, we surveyed the distribution of 24 mRNA regulators and P granule proteins in the P_4_ blastomere (Table 1). At this stage, P granules associate with the surface of the nucleus forming lobes that extend from the nuclear surface into the cytoplasm (Fig. EV5). Most of the proteins in the survey showed enrichment in perinuclear lobes (Fig. EV5). Strikingly, the two ribosomal proteins and five subunits of the 43S pre-initiation complex included in our survey showed the opposite pattern: enrichment in the cytoplasm and depletion from perinuclear lobes (Fig. EV5).

**Table 1.**
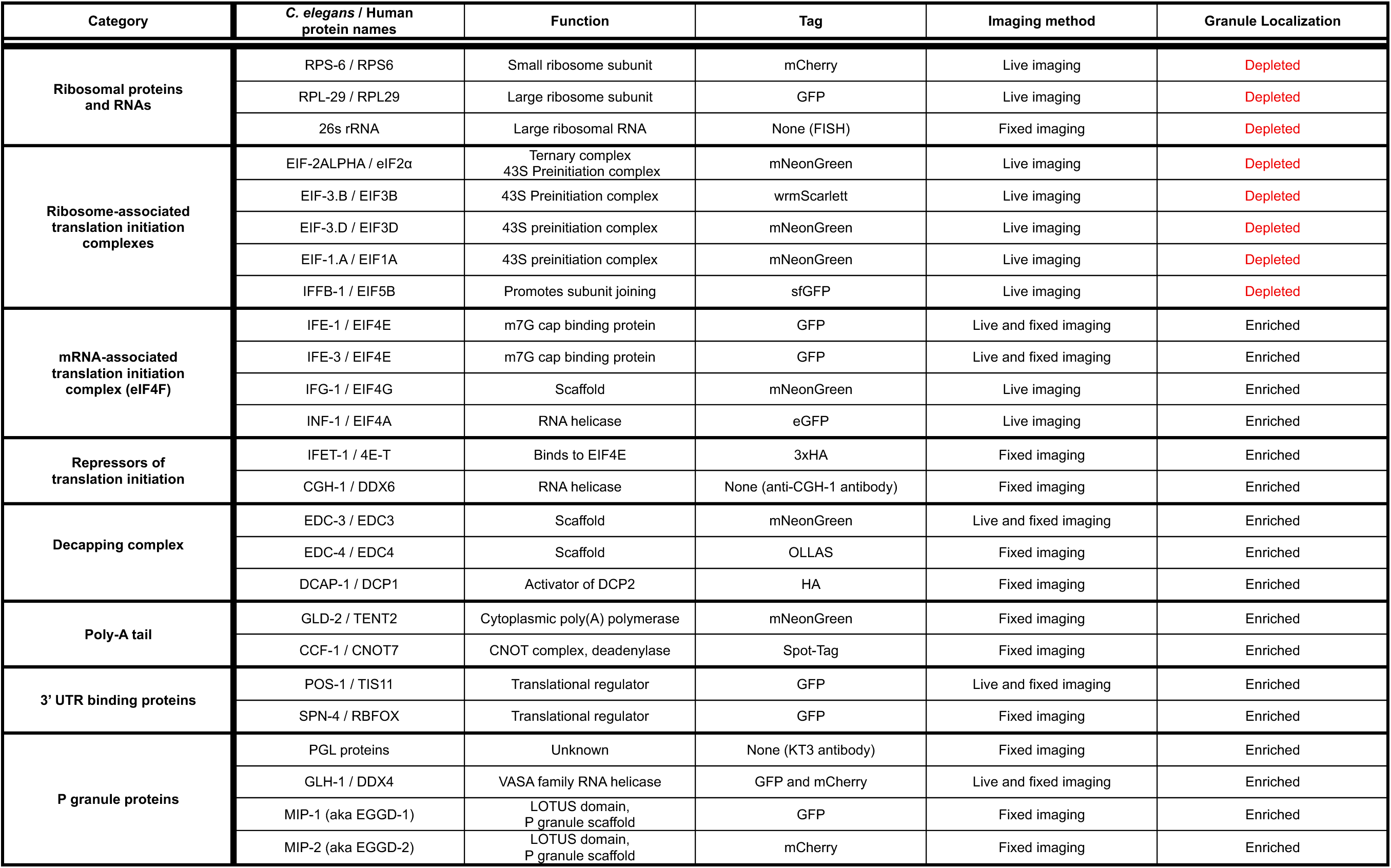

To investigate this pattern further, we compared the intensity in nuclei, P granules and cytoplasm of 26S rRNA (detected by smFISH), the small subunit ribosomal protein RPS-6 fused to mCherry, the ribosome-associated initiation factor EIF-1.A fused to mNeonGreen, and the mRNA cap binding protein IFE-1 fused to GFP (Fig. 5). These analyses confirmed that ribosome-associated factors are depleted from P granules, in contrast to IFE-1 which is enriched in P granules. We observed similar depletion and enrichment patterns for non-perinuclear P granules in P_2_ and P_3_ blastomeres (Fig. EV5). The extent of depletion of ribosome-associated factors from P granules approached that observed in nuclei (Fig. EV5). We conclude that ribosomes and associated initiation factors are depleted from P granules to levels nearing levels in nuclei. In contrast, proteins that interact with mRNAs independent of ribosomes accumulate in P granules.

**Fig. 5:**
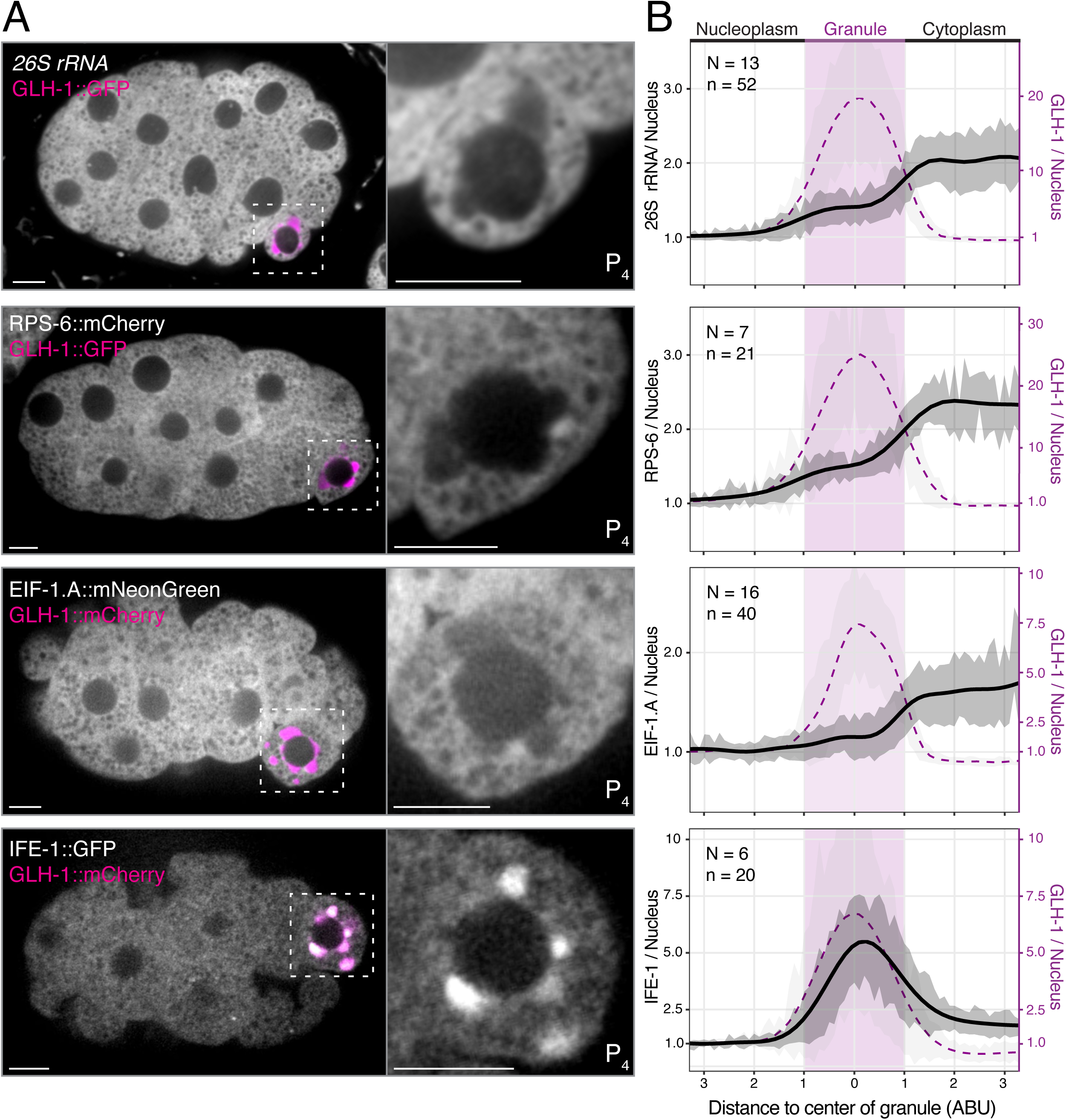
P granules are depleted of ribosomes and 43S pre-initiation complexes. **(A)** Photomicrographs of P_4_-stage embryos expressing a tagged GLH-1 (GFP or mCherry) and either fixed for *in situ* hybridization against 26S rRNA, or imaged live for RPS-6::mCherry, EIF-1.A::mNeonGreen, or IFE-1::GFP expression. Insets show single-plane photomicrographs of the P_4_ blastomere from the embryo shown in A without the GLH-1 channel. Scale bars are 5µm. **(B)** Graphs showing average fluorescence intensities of line scans centered on the local maximum of a GLH-1 granule and extending into the nucleus and cytoplasm. N= # of embryos sampled and n= # of granules analyzed. Each line represents a different GLH-1 granule. The X-axis represents distance normalized to an average GLH-1 granule radius for each dataset (1 radius = 1 unit, Methods). The left Y-axis (black) shows fluorescence signal (arbitrary units) normalized to mean nucleoplasmic signal (y = 1). The black trace represents a generalized additive model fitted curve to the population mean of line scans. The dark gray ribbon represents ± 1 standard deviation of the nuclear normalized line scans calculated across 0.1 radius bins. The right y axis and magenta curve show mean normalized GLH-1 fluorescent signal (arbitrary units) normalized to nucleoplasmic GLH-1 (y = 1). GLH-1 enrichment values varied dependent on the tag used, with GLH-1::GFP concentrating in granules more efficiently than GLH-1::mCherry.

## Discussion

Germ granules concentrate maternal mRNAs in germline precursors and were recently proposed to activate the translation of *nanos* mRNA in *Drosophila* (Chen *et al*, 2024; Ramat *et al*, 2024). We used single molecule techniques to examine the distribution and translation status of P granule-enriched mRNAs in *C. elegans* embryos. We find that P granules concentrate untranslated mRNAs but do not regulate translation, which occurs primarily in the cytoplasm. Together with prior findings, our observations suggest the following working model. Maternal mRNAs are regulated by RNA-binding proteins that distribute between the cytoplasm and P granules, but function independently of P granules. Before translational activation, a subset of untranslated mRNA molecules condenses with the germ plasm protein MEG-3 to help stabilize P granules. Asymmetric segregation of MEG-3 with germ plasm enriches these mRNAs in P blastomeres. At translation activation, mRNAs are translated in the cytoplasm but rarely in granules, which are depleted of ribosomes. We discuss each aspect of this model and its potential relevance to germ granules in other systems.

### P granules are depleted of ribosomes

A main finding of our study is that, compared to surrounding cytoplasm, P granules are depleted of 26S rRNA, ribosomal proteins RPS-6 and RPL-29 and translation initiation factors that associate with the 40S subunit to form the 43S pre-initiation complex (Table 1) (Brito Querido *et al*, 2024). Schisa *et al*, (2001) and Marnik *et al*, (2019) reported that 18S rRNA is excluded from P granules in meiotic germ cells in adult hermaphrodites, raising the possibility that depletion of ribosomes is a characteristic of P granules throughout germ cell development. Marnik *et al*, also reported that immunoprecipitates of the P granule protein GLH-1 are depleted of ribosomal proteins and proteasome subunits. Subunits of the cap-binding complex eIF4F (eIF4E, eIF4A, and eIF4G), in contrast, have been reported to enrich in P granules in both adult germlines and embryos (Table 1) (Huggins *et al*, 2020; Gajjar *et al*, 2025). Using labeled dextrans of varying sizes, Updike *et al*, (2011) observed that embryonic P granules exhibit a size-exclusion barrier between 40 and 70kDa, which would be sufficient to exclude assembled ribosomes (∼3.2 MDa), but theoretically should also exclude eIF4G (∼175 kDa) which enriches in the granules. Taken together, these observations indicate that P granules form a compartment that is permeable to translation initiation complexes assembled on mRNAs but mostly impermeable to translating ribosomes and 43S pre-initiation complexes.

Several observations suggest that germ granules in other organisms may also exclude ribosomes. Westerich et al. (2023) reported that, in zebrafish, large and small subunit ribosomal proteins accumulate at the granule periphery whereas eIF4G and eIF4E concentrate inside germ granules. Electron microscopy studies confirmed enrichment of ribosomes and polysomes in cytoplasm immediately adjacent to the granules. Similarly, electron microscopy studies in *Drosophila* reported polysomes protruding into the cytoplasm from the surface of polar granules (Mahowald, 1968). Chen *et al*, 2024 reported that the ribosomal protein RPS6 enriches in germ granules in *Drosophila*, but it remains possible that assembled ribosomes are excluded from the granule interior since *nanos* translation occurs mostly at the granule surface. Exclusion of ribosomes from the granule interior could cause cytoplasmic ribosomes to cluster at the granule periphery where they can access mRNAs tethered to the granules. We did not observe such surface clustering in *C. elegans*, possibly because P granules are more dynamic than *Drosophila* or zebrafish germ granules and allow mRNAs and regulators to exchange readily with the cytoplasm. Weak ribosome clustering at the granule border might also have been missed by our imaging. It will be important to explore whether ribosome exclusion is a general property of germ granules that is selective and important for function or arises as a secondary consequence of the material properties of the granules.

### The translation competency of mRNAs enriched in P granules is determined by regulators that distribute between P granules and the cytoplasm, but function independently of the granules

A key challenge when defining the roles of condensates is to distinguish functions emergent to the condensates from functions intrinsic to the proteins that concentrate in the condensates (Mateju & Chao, 2022; Putnam *et al*, 2023). In *C. elegans*, germ granules can be genetically uncoupled from other germ plasm components using mutations in MEG-3 and MEG-4. MEG-3 and MEG-4 are intrinsically-disordered proteins that condense with untranslated mRNAs to form low-dynamic clusters that adsorb to the surface of P granules and stabilize the granules in germ plasm (Folkmann *et al*, 2021; Schmidt *et al*, 2021). In the absence of *meg-3 meg-4*, mRNAs are not recruited to the granules and P granule condensates are not stabilized in germ plasm. Germ plasm proteins, including translational regulators like POS-1 that enrich in P granules, however, still segregate asymmetrically to P blastomeres in *meg-3 meg-4* embryos (Wang *et al*, 2014; Cassani & Seydoux, 2022). We reported previously that *nos-2* and *xnd-1* are translated in *meg-3 meg-4* embryos but produce lower protein levels, which we hypothesized was due to lower numbers of mRNA molecules available for translation (Cassani & Seydoux, 2022). Consistent with this hypothesis, we find that the cytoplasmic pool of *nos-2* and *xnd-1* is reduced in *meg-3 meg-4* embryos, but translation onset and efficiency (percent of mRNAs in the cytoplasm engaged in translation) is not reduced relative to wild-type. We conclude that mRNA recruitment by germ granule scaffolds, like MEG-3/4, boost protein production by concentrating maternal mRNAs in germline blastomeres, but do not regulate translation.

If germ granules are dispensable for translational regulation, what activates Nanos translation in P blastomeres? Our observations support a role for RNA-binding proteins that enrich in germ granules but operate independently of the granules. Building on the findings of Jadhav et al., (2008), we show that *nos-2* translation follows an increase in POS-1 levels in P blastomeres relative to the translational repressor SPN-4 and that this increase does not require P granules. POS-1 regulates the poly(A) tail length of hundreds of mRNAs, including *nos-2* and *xnd-1* (Elewa *et al*, 2015). *pos-1* mutants fail to activate *nos-2* translation, and do not complete embryogenesis due to a failure to specify the fates of the C, D and P_4_ blastomeres (Tabara *et al*, 1999; D’Agostino *et al*, 2006; Elewa *et al*, 2015). We speculate that the progressive rise in POS-1 concentration in germ plasm changes the combinatorial landscape of RNA-binding proteins on 3’ UTRs, allowing for temporal regulation of translation across a range of transcripts. POS-1 likely associates with target mRNAs inside and outside the granules, but mRNAs are translated most efficiently outside the granules where ribosomes are most abundant.

Translation of germ granule-associated mRNAs in *Drosophila* is also temporally regulated, with different granule transcripts translated at different embryonic stages (Rangan *et al*, 2009). The observation that *nanos* transcripts tagged with SunTag repeats are translated at the surface of germ granules with their 5’ end oriented towards the cytoplasm suggested that the granules regulate translation by segregating translational repressors and activators to distinct internal and surface compartments (Chen et al 2024, Ramat et al 2024). Consistent with this hypothesis, mutations in granule scaffolds that disrupt germ granule composition or architecture without affecting *nanos* mRNA recruitment to the granules prevent *nanos* translation. An alternative hypothesis is that the granules activate Nanos translation indirectly by co-localizing *nanos* mRNA with its regulators, but not by regulating their activity.

### mRNAs in P granules are eventually turned over in the germline founder cell

Several studies reported that *nos-2* RNA enrichment in granules becomes reduced in the germline founder cell P_4_ compared to other P blastomeres (Lee *et al*, 2020; Parker *et al*, 2020). This observation has been interpreted as relocalization of *nos-2* mRNA from granule to cytoplasm during translation. Our new findings challenge that interpretation by demonstrating that mRNA levels in P granules decrease without a compensatory increase in the cytoplasm during the lifetime of the P_4_ blastomere. The simplest explanation for this observation is that the granule pool of *nos-2* and other maternal mRNAs is turned over, either within the granules or after release to the cytoplasm. Consistent with the former possibility, in *Drosophila*, germ granules have been shown to recruit decay factors and become sites of mRNA decay after pole cell formation (Hakes & Gavis, 2023). Together these observations suggest that the pool of mRNAs associated with germ granules is not maintained after the granules have completed their segregation into the PGCs.

### A unifying model for germ granule function

If germ granules are not required to activate mRNA translation and eventually lose associated mRNAs, why do mRNAs localize to germ granules? We have suggested that the primary function of germ granules is to transmit Argonautes and other small RNA machinery to the PGCs (Ouyang *et al*, 2019) and that mRNAs are recruited to the granules to scaffold and stabilize the granules (Scholl *et al*, 2024). MEG-3 requires RNA *in vivo* and *in vitro* to condense into clusters that adsorb to the surface of P granules and lower surface tension (Schmidt *et al*, 2021; Folkmann *et al*, 2021; Putnam *et al*, 2019). We speculate that the scaffolding function of mRNAs is only required in oocytes and early embryos where germ granule proteins are relatively dilute and require mRNAs to enhance their condensation in germ plasm. Once the granules have segregated into the much smaller, hyper-concentrated environment of the germline precursors, mRNAs are no longer needed and can be turned over as these cells prepare for the maternal-to-zygotic transition. Our observations also suggest that, for a subset of mRNAs (e.g. *nos-2 and xnd-1)*, binding to germ granule scaffolds, such as MEG-3/4, increases the pool available for translation in germline precursors, which indirectly boosts protein production. MEG-3 distributes between the granules and a cytoplasmic gradient that segregates to P blastomeres (Wang et al., 2014) and so theoretically could bias the segregation of, not only P granule-associated mRNAs, but also mRNAs in the cytoplasm. Alternatively, mRNAs in P granules could redistribute from the granules to the cytoplasm after cell division.

*zif-1* did not enrich in the P_4_ cytoplasm as robustly as *nos-2* and *xnd-1*, indicating that MEG-3/4 do not impact the cytoplasmic concentration of all P granule-enriched transcripts to the same extent. We suggest that, while most mRNAs bind to germ granule scaffolds to stabilize the granules, a subset uses this association to increase their own concentration in germline precursors. Whether these mRNAs are translated at the granule surface or away from the granules likely depends on the condensation dynamics of the germ granule scaffolds. We suggest that, during the syncytial stages of *Drosophila* embryogenesis, germ granule scaffolds and associated mRNAs must remain highly condensed in the granules to avoid dilution into somatic cytoplasm, thereby restricting translation to the granule surface. In contrast, in embryos that cellularize early like *C. elegans,* germ granule scaffolds distribute between the granules and cytoplasm, allowing for translation of associated mRNAs in the cytoplasm. In both cases, translation is controlled by factors that concentrate in the granules, but don’t require the granules to modulate activity.

### Limitations of this study

Our conclusions rest on measurements in fixed embryos, which capture snapshots of mRNA and protein distribution rather than the behavior of individual molecules over time. Because we have not examined RNA dynamics directly, we do not know whether mRNAs exchange between the granules and the cytoplasm. Because nascent-peptide detection has a sensitivity threshold, it is possible that we missed mRNAs translated by single ribosomes. The low translation frequencies we observe in P granules should be interpreted as an upper threshold for translational exclusion and do not imply a complete absence of translation. We inferred ribosome depletion from a limited panel of ribosomal proteins, initiation factors and rRNA, rather than from direct visualization of assembled ribosomes and/or polysomes and therefore cannot exclude the possibility that some ribosomes access the granule interior and/or cluster at the surface.

## Materials and Methods

### *C. elegans* strains and maintenance

*C. elegans* strains were maintained at 20°C on NNGM plates (IPM Scientific Cat#11006-548) seeded with OP50 *E. coli* (CGC) according to standard methods(Brenner, 1974), unless otherwise stated. Strains used in this study are listed in the strain table (Table S1).

### RNAi

RNAi knockdowns were performed by feeding as previously described (Timmons & Fire, 1998). RNAi plasmid vectors were miniprepped (Qiagen Cat # 27104) from clones drawn from the Ahringer RNAi Library (Kamath *et al*, 2003, originally obtained from Source Bioscience) according to kit instructions. Purified RNAi vectors were transformed into competent HT115 *E. coli* (CGC), grown for 5-7 hours at 37°C in LB broth (Apex Bioresearch Products) with ampicillin (100 µg/mL, Sigma Aldrich). Cultures were then induced with 1 mM IPTG (Gold Biotechnology) for 30 minutes at 37°C, concentrated by centrifugation, resuspended, and seeded on RNAi plates (NNGM + 50 µg/ml carbenicillin+1 mM IPTG, IPM Scientific). Each transformation of any RNAi vector was accompanied by a transformation of the empty vector backbone plasmid, L4440 (Gift of Andrew Fire, Addgene plasmid # 1654) to serve as a vector-only control in subsequent RNAi experiments. Seeded RNAi plates were allowed to grow at room temperature overnight before use. L4 hermaphrodites were transferred onto seeded RNAi plates and allowed to grow overnight to young adulthood. Empty vector controls were performed under the same conditions as the RNAi treatment.

In experiments using combined *meg-1/meg-2(RNAi)* or *meg-3/meg-4(RNAi)*, RNAi cultures of each vector were grown separately and mixed 50/50 by volume before seeding. Worms were grown at 25°C overnight before being transferred to 20°C for at least 1 hour prior to dissection to equilibrate (Cassani et al 2022). In the cases where *meg-3(ax3055) meg-4(ax3052)* animals were compared to *meg-1(vr10) meg-2(RNAi)* treated animals, both the wild-type and *meg-3 meg-4* animals were grown on empty vector bacteria under the same conditions as *meg-1(vr10) meg-2(RNAi)* treated animals.

In *meg-1(vr10) meg-2(RNAi) + spn-4(RNAi)* experiments, L4 *meg-1(vr10)* hermaphrodites were transferred to control or *meg-2(RNAi)* plates overnight at 25°C before being transferred to either empty vector or *spn-4(RNAi)* plates for 8 hours at room temperature before collection.

### smFISH probes

smFISH probes used in this study were designed using the Stellaris Probe Designer (Biosearch Technologies) and either ordered synthesized by Biosearch Technologies or synthesized in house according to the protocol outlined by Scholl et al(2024) adapted from Gaspar et al. (2018) using oligo pools ordered from IDT and either Cy3 (Lumiprobe Cat#11020) or Cy5 (Lumiprobe Cat#13020) NHS esters. To visualize RNA-rich granules, including P granules and germline P-bodies, we used a probe complementary to the splice-leader SL1 sequence labeled with fluorescein (Sequence: CTCAAACTTGGGTAATTAAACC, Biosearch Technologies) to generate granule masks (see Quantification of translating mRNAs).

### Single-Molecule FISH (smFISH)

Gravid adults were dissected to extrude embryos onto poly-L-lysine-coated slides in 1x egg buffer with Tween 20 (25 mM HEPES, 118 mM NaCl, 48 mM KCl, 2 mM EDTA, 5 mM EGTA, 0.1%Tween 20, 10 mM levamisole), or M9 buffer. Embryos were freeze-cracked on dry-ice-chilled metal blocks and immediately submerged in -20°C methanol for a minimum of 1 hour to dehydrate. Samples were subsequently rehydrated through a 1:1 mixture of cold methanol and PBS + 0.1%Tween 20 (PBSTw) followed by five washes in 100% PBSTw. Samples were fixed for one-hour in 4% paraformaldehyde in PBS at room temperature in a humid chamber and washed sequentially four times in PBSTw, twice in 2x SSC, once in wash buffer (10% formamide in 2x SSC) and finally equilibrated in hybridization buffer (10% formamide, 2x SSC, 200 µg/mL Ultrapure BSA (Applied Biosystems AM2618), 2 mM Ribonucleoside Vanadyl complex (NEB S1402S), 0.2 mg/mL yeast total RNA, 10% dextran sulfate Leuconostoc spp, MW >500,000 (Sigma D8906) for at least 30 minutes at 37°C. Hybridization was performed overnight at 37°C in sealed coverslip chambers using 80–100 µL of hybridization buffer containing smFISH probes at 5-10µM. Post-hybridization, coverslips were removed and samples were washed twice in wash buffer at 37°C for up to 30 minutes each. Samples were washed twice in 2xSSC, once in PBSTw, twice in PBS and mounted in either Vectashield PLUS with DAPI (Vector Laboratories H-1900) and sealed or Vectashield Vibrance (Vector Laboratories H-1800) and allowed to cure overnight before imaging.

### Immunofluorescence

Embryos were dissected in egg buffer or M9 onto poly-L-lysine-coated slides, freeze-cracked, and immediately fixed in -20°C methanol for a minimum of 20 minutes. Slides were blocked in PBS containing 0.1% v/v Tween 20 and 0.1% w/v BSA (blocking buffer) for 30 minutes at room temperature, then incubated with primary antibodies diluted in blocking buffer overnight at 4°C to minimize background signal. Following three 10-minute washes in blocking buffer, samples were incubated with secondary antibodies for 1-2 hours at room temperature in the dark.

Nanobodies were added with secondary antibody incubations when used. After a final series of three 10-minute washes in blocking buffer, slides were mounted using Vectashield PLUS with DAPI and sealed. Primary antibody concentrations were as follows: Rat anti-OLLAS L2 1:100 (Novus #06713, gift of Jeremy Nathans), Rabbit anti-CGH-1 1:1000 (Alessi *et al*, 2015, Gift of John Kim), Rabbit anti-HA-Tag 1:200 (Cell Signaling #3724), Spot-Label Alexa Fluor 568 nanobody 1:100 (Proteintech #ebAF568), Mouse KT3 (anti-P granule) 1:1000 (Takeda *et al*, 2008, DSHB). Secondary antibodies included Alexa Fluor® 647 Goat Anti-Rabbit IgG (H+L) (Thermo Fisher, A-21245) 1:200, Alexa Fluor® 568 goat anti-Rat IgG (H+L) (Thermo Fisher, A-21245) 1:200 and Goat Anti-Mouse IgA alpha chain DyLight® 550 (Abcam, ab97012).

### Simultaneous Immunofluorescence and smFISH

#### Sample Preparation and Fixation

Embryos were dissected onto the middle well of a three-well slide coated with poly-L-lysine in egg buffer, freeze-cracked, and submerged in -20°C methanol for a minimum of 20 minutes. Samples were rehydrated through a 1:1 mixture of 1x PBS and methanol into 1x PBS, followed by a brief rinse in PBSM (1x PBS + 5 mM MgCl_2_). Embryos were then fixed in 4% paraformaldehyde in PBSM for 10 minutes at room temperature.

#### Permeabilization and Pre-hybridization

Following fixation and PBSM washes, samples were permeabilized for 10 minutes at room temperature in permeabilization buffer (PBSM, 0.1% Triton X-100, 5 mg/mL BSA, 10 U/mL SUPERase·In (ThermoFisher AM2694), 2 mM Ribonucleoside Vanadyl Complex (NEB S1402S, RVC). Slides were washed in PBSM and equilibrated in pre-hybridization buffer (10% formamide, 2x SSC, 5 mg/mL BSA, 10 U/mL SUPERase·In, and 2 mM RVC) for 30 minutes at room temperature.

#### Hybridization

Simultaneous primary antibody binding and probe hybridization was performed by incubating slides for 3 hours with smFISH probes and primary antibody or nanobody in hybridization buffer (10% formamide, 2x SSC, 0.2 mg/mL Ultrapure BSA, 10% dextran sulfate(Sigma D8906), 1 mg/mL yeast RNA, 10 U/mL SUPERase·In, 2 mM RVC) at 37°C in a humid chamber with samples protected by coverslips. smFISH probes were diluted to 200 nM. Anti-HA primary antibody (Cell Signaling) was diluted to 10 µg/mL. Anti-ALFA FluoTag-X2 nanobody (NanoTag Biotechnologies, N1502) was used at a 1:200 dilution.

#### Secondary Detection and Mounting

Coverslips were removed, and samples were washed three times for 5 minutes each in pre-warmed 37°C wash buffer (10% formamide, 2x SSC). Slides were then incubated with Alexa Fluor® 647 Goat Anti-Rabbit IgG (H+L) diluted in wash buffer at 1:2000 for two consecutive 20-minute periods at 37°C. Following three final 10-minute washes in 37°C 1x PBS, samples were mounted using Vectashield Vibrance with DAPI and sealed.

### Confocal microscopy

#### Fixed imaging

Z-stack images of embryos were acquired using an inverted Zeiss Axio Observer equipped with an iXon Life 888 EMCCD camera (Andor) or an upright Zeiss Axio Imager Z1 equipped with an ORCA-Fusion BT digital CMOS camera (Hamamatsu). Both microscopes use CSU-W1 SoRa spinning disk scan heads (Yokogawa), 63x oil objective (Zeiss, 1.4 NA), 2.8× relay lens (Yokogawa), and are controlled by Slidebook software (Intelligent Imaging Innovations). All Z-stack images were acquired using a 0.27 μm step size. Exposure times and power settings were dependent on the experiment but were kept consistent between replicates of the same probes and antibodies.

#### Live imaging

Embryos were dissected from young adult hermaphrodites onto 22 x 22 mm coverslips in M9 buffer. Dissected embryos were mounted on 2% agarose pads or in M9 buffer containing 20 µm polystyrene beads (Bangs Laboratories, PS07003) and sealed with Vaseline for live imaging. Confocal microscopy was performed using an inverted Zeiss Axio Observer equipped with a CSU-W1 SoRa spinning disk scan head (Yokogawa) and an iXon Life 888 EMCCD camera (Andor) using a 63× objective with a 2.8× relay lens (Yokogawa) capturing ∼7-10 µm Z-stacks at 0.4 µm to 0.5 µm step sizes. Green and red fluorescent channels were acquired sequentially for each plane using 100-150ms exposure times to capture dynamic protein distributions across P blastomeres.

### Quantification of mRNA spots and translating mRNAs

#### Image Pre-processing and Cell Isolation

Raw images were pre-processed using custom Jython scripts in Fiji. Images underwent channel-specific Gaussian blurring (σ=1 for RNA and protein channels, σ=2 for granule markers) and rolling-ball background subtraction (30-pixel radius) followed by a linear intensity correction for the signal decay in deeper Z-slices based on a Z-axis profile of a user-defined region of embryo tissue. When needed, nuclear background was removed by generating a DAPI mask and applying a Gaussian-smoothed fill. P-lineage blastomeres were isolated for quantification through manual 3D region-of-interest (ROI) cropping and interpolation across Z-slices. These ROI sets were also used to approximate the volume of the P_4_ blastomere (total measured ROI area multiplied by step size).

#### Spot Detection and 3D Colocalization

Single-molecule spot detection was executed using the TrackMate plugin in Fiji (Ershov *et al*, 2022)equipped with a Laplacian of Gaussian (LoG) detector. Spots were linked across adjacent Z-planes using a strict maximum linking distance of 0.15 µm in the XY plane to prevent double counting. Under the imaging condition of 0.27 μm/z-step, spots that could not be linked across at least 2 slices were considered non-specific noise and were discarded. To accurately size and classify the initial TrackMate detections, a custom diameter-examining algorithm evaluated each coordinate to determine the optimal diameter of spots that provides the strongest LoG response. The 3D intensity measured from this derived size subsequently served as a classification gate to categorize each signal as either a single transcript or a dense multi-RNA cluster prior to colocalization.

To identify actively translating mRNAs, 3D colocalization between the RNA and protein channels was performed by screening for RNA-protein spot pairs with a x-y distance smaller than a 0.20 μm threshold and x-z distance smaller than 0.40 μm. To correct for the unidirectional spatial off-set often observed between 2 channels, either multi-spectral beads (Invitrogen #T7279) or dual color smFISH samples applying 2 probe sets of different colors with the same sequences can be imaged as a control on the same day of experiment with the same imaging conditions. The spatial offset (dX, dY, dZ) is calculated from the control. An opposite vector is then applied to other images of the same experiments to correct the offset.

#### Granule Spatial Mapping

To determine the spatial positioning of RNAs relative to germ granules, binary granule masks were generated manually by adjusting the threshold to ensure that the majority of SL1 signal above cytoplasmic background was captured. Next, 3D Euclidean Distance Transform (EDT) maps were computationally generated from these binary masks of the thresholded granule channel. Individual RNA spots and colocalized translation pairs were then mathematically mapped against these EDT volumes to classify them as "Inside" or "Outside" of the condensate, and to extract their absolute distance to the nearest condensate surface. To account for the resolution limits of our imaging system, we considered any RNA or translation spot within 150 nm of a defined granule surface to be granule-associated and counted it as “inside”.

#### RNA Quantification

Raw fluorescent intensities were converted to absolute mRNA copy numbers using a custom R pipeline. The algorithm isolated the single-molecule RNA population by applying a specific size gate (excluding clusters ≥ 0.4 µm in diameter) and then utilized a 95% trim, discarding the brightest 5% of isolated spots, to fit a Gaussian distribution and calculate the mean "unit" intensity of a single transcript. This calibrated single-transcript intensity was used to calculate the total number of mRNAs in multi-RNA clusters by dividing the total intensity of the cluster by the calibrated single transcript intensity value. This corrected mRNA count was used to calculate an absolute number of RNAs in total, inside, and outside of granules.

#### Specific considerations for quantification of nos-2 translation sites

Because *nos-2* mRNA and mature NOS-2 protein accumulate on P granules, RNA spots in granules that colocalize with protein spots could represent either genuine translation sites or colocalization of mRNA and protein post-translation. To avoid over-counting translation spots in granules, we took the following measures. First, we only scored early P_3_ blastomeres where the anti-HA signal was above background but not yet entirely coincident with the SL1 granule pattern (as is typical in later P_3_ and in P_4_ blastomeres). Second, each anti-HA spot co-localized with mRNA in a granule was counted as one translating mRNA molecule, even if the spots overlapped an RNA cluster. Anti-HA spots colocalized with RNA in granules were not any brighter than anti-HA spots colocalized with RNA in the cytoplasm, consistent with only one translation event.

#### Quantification of mRNA alone

To quantify mRNA spots without protein staining, the same logic used above to count RNAs was duplicated to allow for simultaneous quantification of RNA in two fluorescent channels. All steps of the pipeline remained the same, but colocalization between the two target channels was discarded.

Custom Fiji/Python scripts used in the pipeline were developed with assistance from GPT-4o and o3 (OpenAI) and Gemini 3.0 Flash (Gemini). All AI generated code was subjected to manual review and output validation before use in final figures.

### Quantification of bulk NOS-2 protein signal in P blastomeres

Images were processed using ImageJ/Fiji. A Gaussian blur of sigma = 1 was applied before applying rolling ball background subtractions of radius = 50 on all images prior to quantification. The slice closest to the center of the P blastomere was identified, and a 10-slice (2.7µm) sum projection was made with that slice at its center. The P-blastomere boundary was manually traced to create an ROI and the mean NOS-2 signal intensity was measured. Then, the same ROI was moved to a somatic region of the same embryo and measured there to serve as background. Final values were calculated by subtracting the somatic mean value from the P blastomere mean value. Data were analyzed and plots were generated in R, and the final plots were formatted in Adobe Illustrator.

### Quantification of fixed GFP::POS-1 and SPN-4::GFP signal in P blastomeres

Images were processed in ImageJ/Fiji. The center of the P-blastomere was identified based on DAPI signal and the cytoplasm of the P blastomere was traced outlineds so as to exclude thenucleus. The resulting ROI was measured to find the mean signal of the P blastomere cytoplasm in that plane. Because a uniform somatic background did not exist for all embryonic stages imaged, a global background signal was determined by comparing somatic means of at least five late-stage embryos which did not contain either protein. This global value was then subtracted from the measured mean P blastomere values to correct for autofluorescence background. Data were analyzed and plots were generated in R, and the final plots were formatted in Adobe Illustrator.

### Line scan quantification of ribosomal factors and translation factors in P granules

#### P_4_ Blastomeres

Using Fiji, 4-µm straight line scans of width 3 pixels were drawn originating in nuclei so that the approximate center of the line scan (2µm) passed through the center of a germ granule and extended into the neighboring cytoplasm, avoiding areas devoid of signal. The raw signal for the GLH-1 channel and target factor was recorded along the trace and collected in .csv files using custom Fiji/python macros. Every granule measured was only sampled once. For each embryo used, a region just outside of the embryo itself was measured in every Z-plane to establish a camera noise background which was then mapped back to the plane each trace was collected from and subtracted from the raw data as a baseline background correction. A floor threshold of 0.1 was used to prevent negative values.

x-axis dimensions were scaled to the Full Width at Half Maximum (FWHM) of GLH-1 signal; the x-axis unit (radius at FWHM) was defined as the maximum absolute distance from the peak center where the background-subtracted GLH-1 intensity remained above 50% of maximum in that granule trace. This average radius at FWHM was designated as 1 unit on the x-axis to allow for a uniform definition of granule boundaries for plotting.

To account for variation in signal intensity between traces, both GLH-1 and target signals were normalized to the baseline nucleoplasmic signal within each trace. This signal was defined as the average signal across the first 0.25µm of the trace, which was always exclusively nuclear. The background subtracted signal across the trace was then divided by this value to anchor the nuclear region at 1. Traces were then spatially normalized by binning X-coordinates into 0.1 relative radius unit bins.

Mean population trends were modeled by fitting thin-plate regression splines via Generalized Additive Models (GAM) utilizing Generalized Cross-Validation (GCV) for automatic smoothing parameter estimation. To evaluate the variance of the pooled population, standard deviations (+/- 1 SD) were calculated independently for each discrete spatial bin across all quantified granules. Statistical processing, grid searches for basis dimension saturation (k) used when fitting GAM models, and dual-axis data visualizations were performed in R utilizing the mgcv and tidyverse libraries.

#### P_3_ Blastomeres

Data collection and processing were identical to the methods used for P_4_ blastomeres. However nuclear baselines were determined on a population basis due to each measured granule being surrounded entirely by cytoplasm. To do this, equally sized ROIs were measured in the center of nuclei for at least five embryos and the average camera-background subtracted signal (as defined previously for P_4_ quantifications) from those ROIs was used to establish a static nuclear baseline for normalization across all granules measured for each target factor.

Custom Fiji macros for automating recording of manually drawn line scans and the downstream computational processing of the resulting datasets for both P blastomere stages were developed and optimized with assistance from Gemini 3.0 Flash (Google). All final scripts, operations, and data extraction pipelines were manually verified, edited and validated against raw datasets by the authors before generating final plots.

### Heat shock of C. *elegans* embryos

Poly-L-lysine-coated glass slides, dissection buffer with 2 mM Levamisole in L-15 media (Gibco, 21083027), and 60 mm NNGM (nematode growth medium) plates were pre-warmed to 32 ℃ in a slide incubator (Boekel Scientific, Slide Moat 240000) on the bench next to a dissection scope. About 40 gravid adult worms were picked together with a small amount of OP50 E. *coli* onto each pre-warmed plate and put back in the slide incubator for 30 min. After heat shock, the worms were cut open in a droplet of ∼10 μL pre-warmed dissection buffer on a pre-warmed poly-L-lysine coated slide to release embryos. The samples were then freeze-cracked and fixed for downstream processing. For negative controls, worms were maintained and dissected at room temperature (∼20 °C).

### CRISPR genome editing

*nos-2* and *zif-1* loci were tagged with 7xHA repeats after the start codon using a CRISPR/Cas9 similar to the method described in Ghanta *et al*, (2021). Briefly, HA repeats flanked by 35 nt homology arms were PCR amplified from a gBlock DNA Fragment (IDT) using Phusion polymerase, purified using AMPure XP beads (NEB) and denatured/renatured according to (Ghanta et al., 2021). Injection mixtures containing 400 ng pRF4::*rol-6(su1006)* plasmid (purified using PureLink HiPure plasmid miniprep kit (Invitrogen, Cat#K210003)), 30 pmol Cas9 (IDT, Cat#1081058), 90 pmol tracrRNA (IDT, Cat#1072532), 95 pmol crRNA, 350 ng repair template were injected into the gonads of strain EGD717, which expresses a GFP::HA Frankenbody transgene. Tagged NOS-2 and ZIF-1 alleles were outcrossed to N2 worms to remove the GFP::HA Frankenbody transgene.

The *xnd-1* locus was tagged by an array of 8 repeats of AlfaTag (8xALFA) at the N-terminus after the start codon following previous CRISPR methods (Paix et al, 2017). Briefly, 8xALFA insert sequences were synthesized as gBlock DNA fragments (IDT). PCR primers with 35-nt overhang (IDT) homologous to the N-terminus of endogenous *xnd-1* were used to generate dsDNA repair template by PCR amplification (Phusion-HF Master Mix, ThermoFisher Scientific, F531L). Roughly 400-800 μL of total PCR reactions were gel purified through a single column (Takara, 740609.5) to produce 10 µL of eluate at a concentration of approximately 1.0-1.5 μg/ μL. This eluate was subsequently mixed with purified Cas9, crRNA and tracrRNA together with reagents targeting dpy-10 as a co-CRISPR marker and injected into the gonads of young adult N2 worms (Paix et al, 2017). 4-5 days post injection, roller F1 worms were singled from jackpot plates and genotyped for the expected insertion. Guide sequences and repair template sequences are available in Table S2.

### Use of AI tools

Claude (Anthropic) was used to generate code in R for data visualization and for language editing. Gemini models (Google) were used for assistance in developing custom Fiji macros, developing quantification scripts in R and Python, and for refining plotting code in R. OpenAI models were used for assistance in developing custom Fiji macros, developing quantification scripts in R and Python. All AI tool output was supervised, vetted, and edited by the authors before final use.

## Acknowledgements.

We thank Bin Wu for sharing protocols and the Seydoux lab for comments on the manuscript. Some strains were obtained from the Caenorhabditis Genetics Center (P40 OD010440).

**The authors declare that they have no conflict of interest.**

## Data and Resource Availability

Source data for all figures are available in S3. Python codes used for image analyses are available on Github.

---

**Figure EV1:**
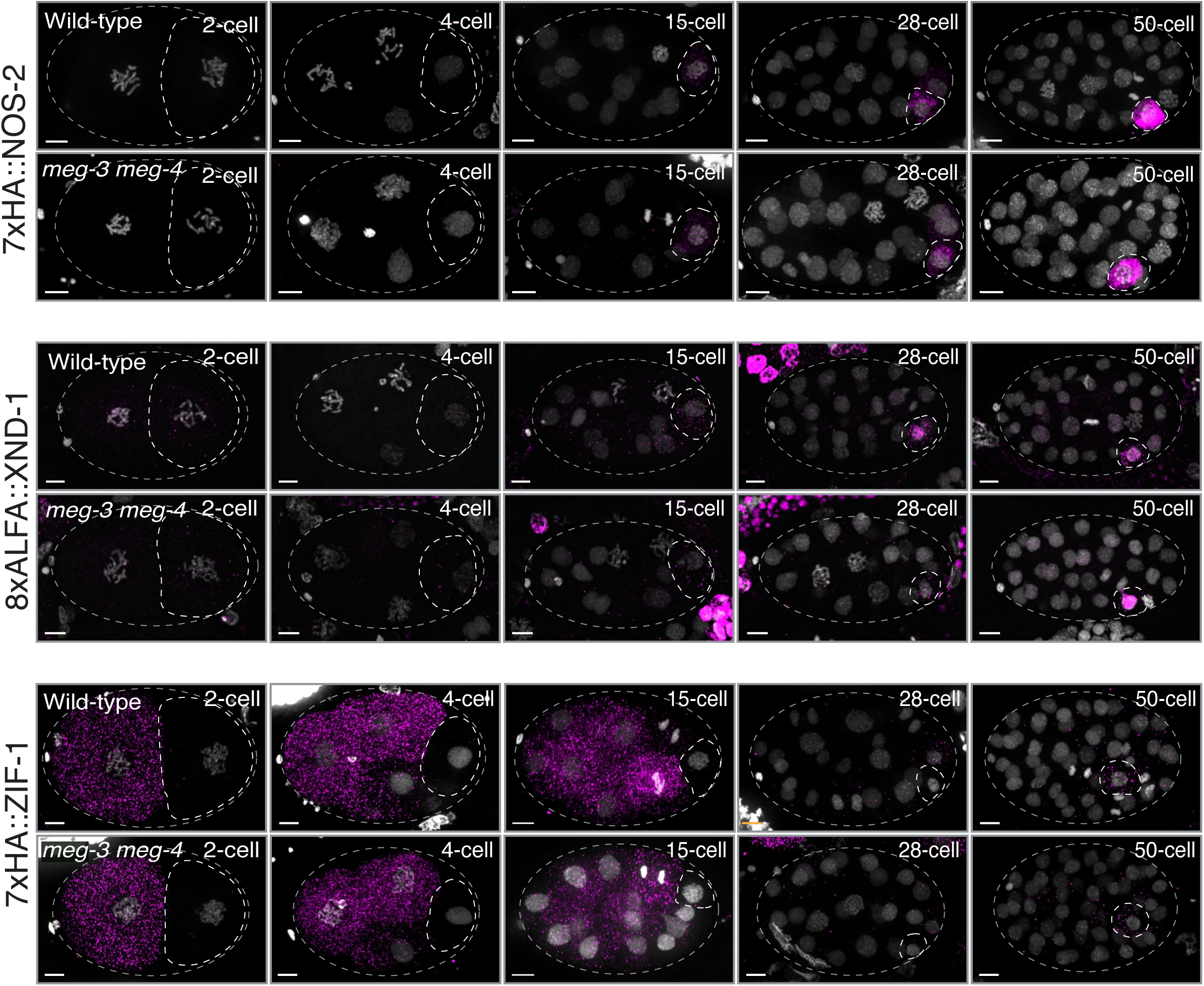
**(A)** Photomicrographs showing maximum projections of embryos immuno-stained for the fusion proteins indicated comparing wildtype and *meg-3(ax3055) meg-4(ax3052)* mutants. Grey dashed lines outline each embryo, and white dashed lines outline P blastomeres. Scale bars are 5µm. Refer to Fig. S1C for lineages showing wild-type expression patterns.

**Figure EV2:**
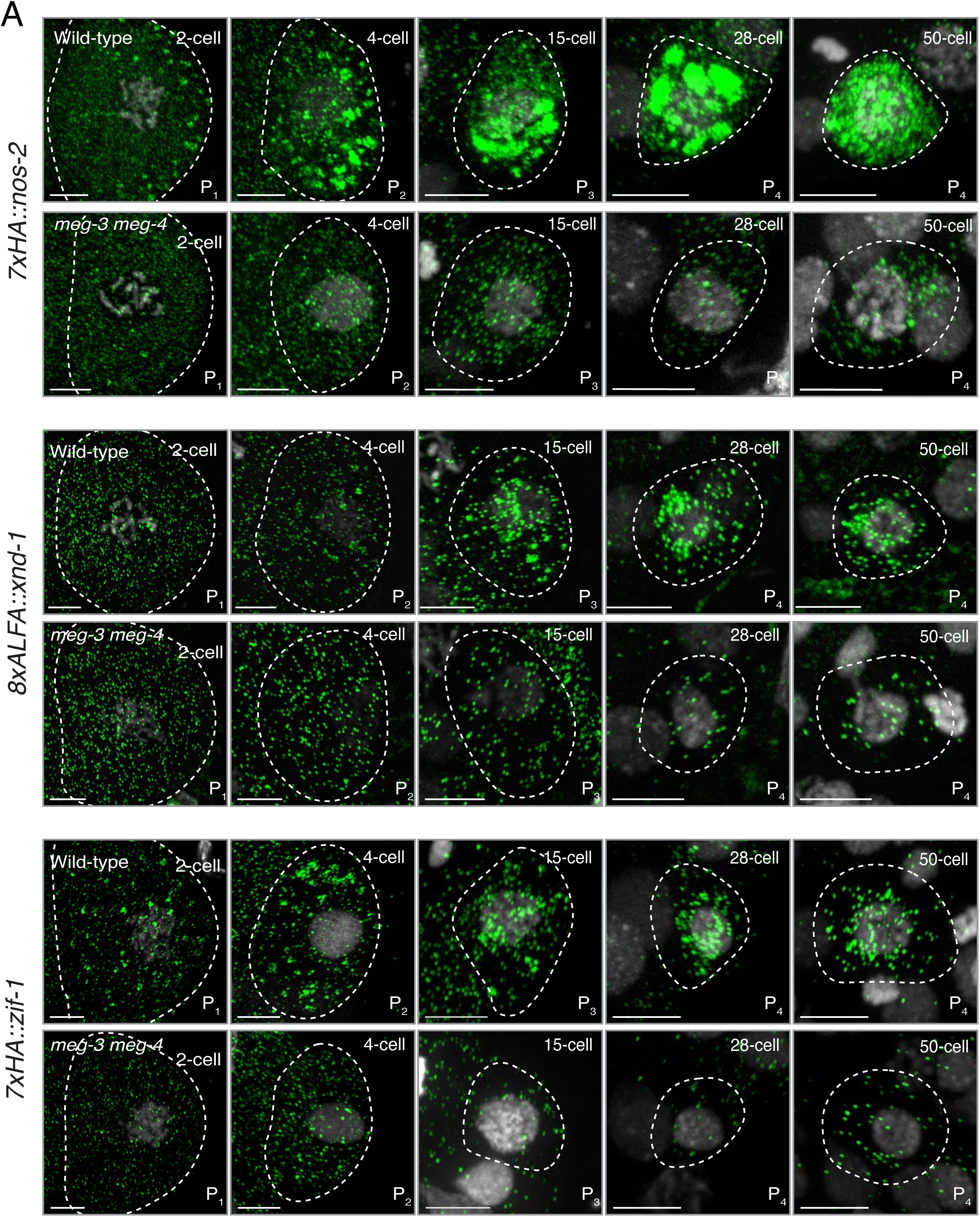
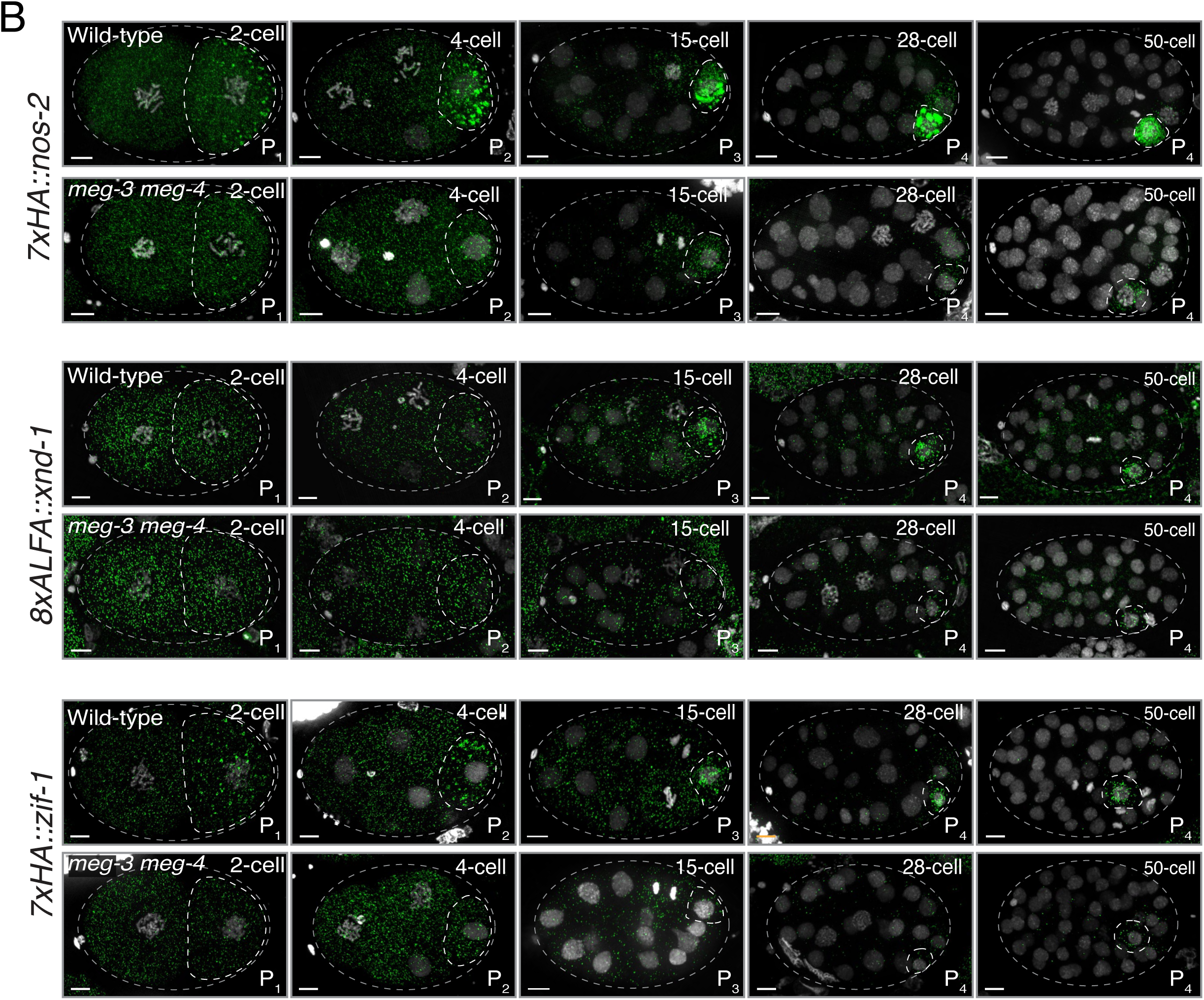
**(A)** Photomicrographs of P blastomeres processed for smFISH against the indicated mRNAs comparing wild-type and *meg-3(ax3055) meg-4(ax3052)* embryos. Dashed lines represent cell boundaries. Images are scaled to accommodate the diminishing sizes of P_1_ through P_4_. Scale bars are 5µm. **(B)** Photomicrographs showing maximum projections of embryos processed for smFISH against the indicated mRNAs comparing wild-type and *meg-3(ax3055) meg-4(ax3052)* embryos. Grey dashed lines outline each embryo and the white dashed lines outline each P blastomere. Scale bars are 5µm.

**Figure EV3:**
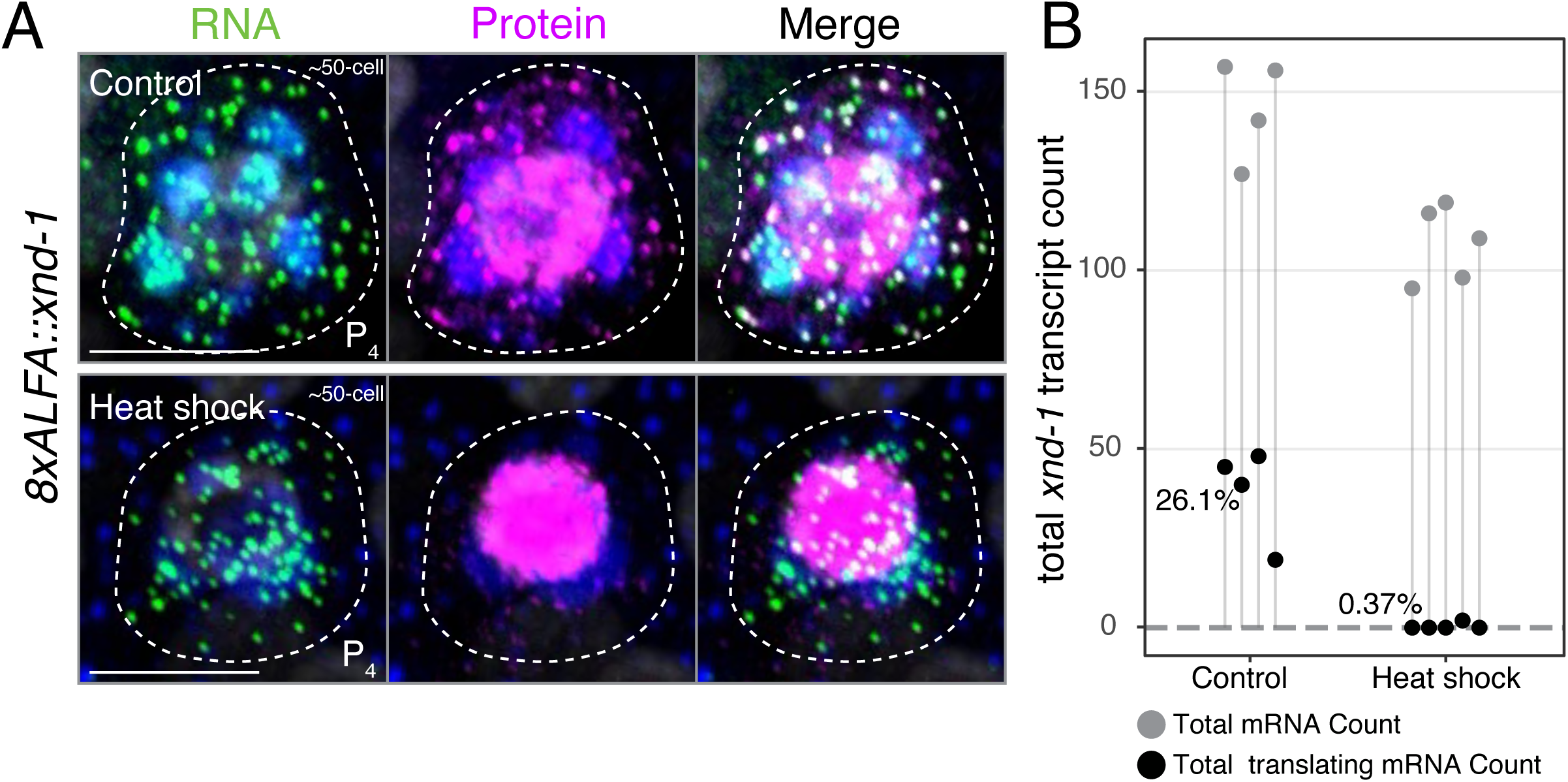
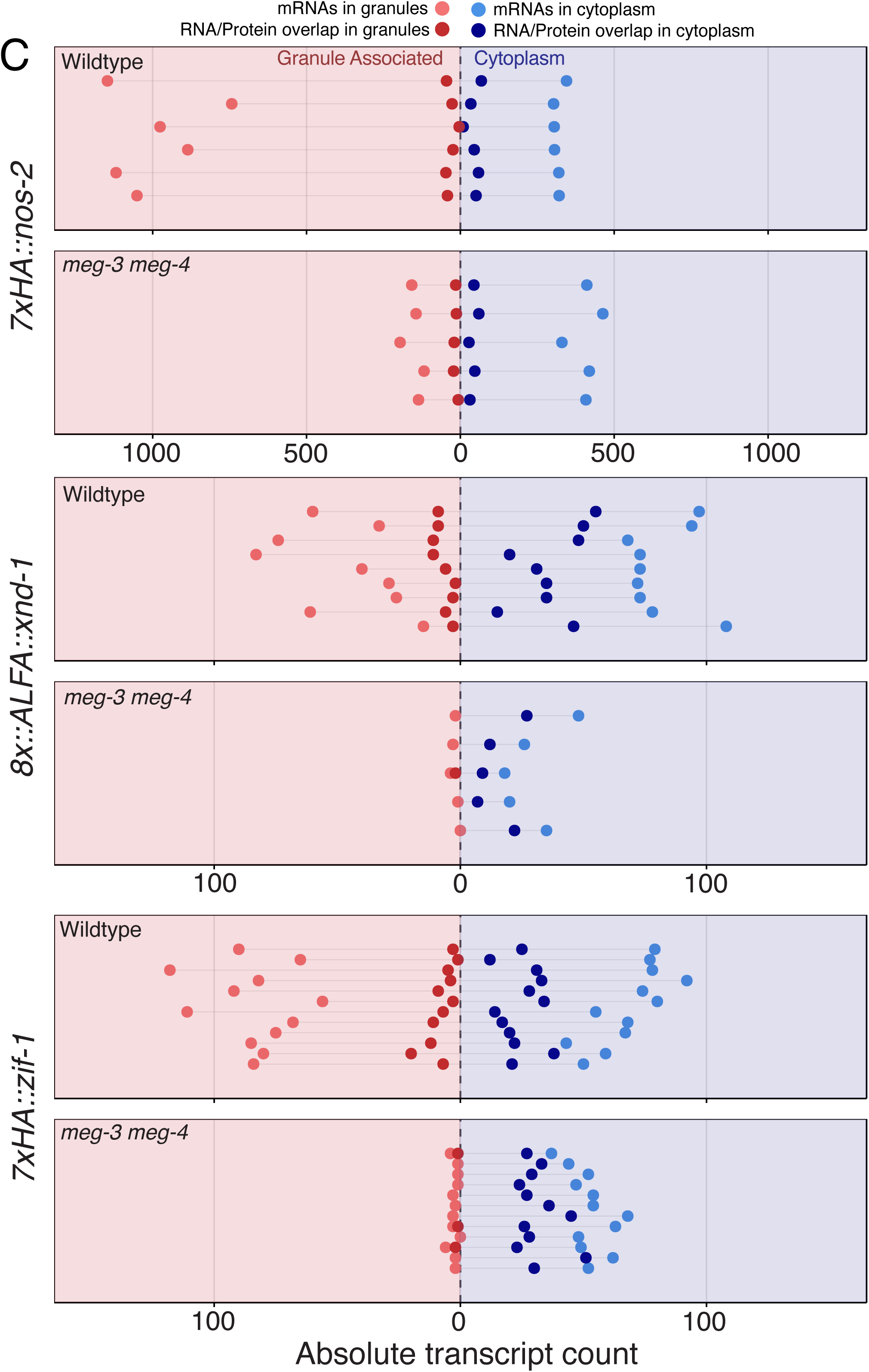
**(A)** Photomicrographs of maximum projections through P_4_ blastomeres expressing *8xALFA::xnd-1* and exposed to either control conditions or 30 minute heat shock prior to fixation and processed as in Fig. 3. Green: RNA, Protein: Magenta, Blue: Granules (SL1), White: DNA. Dashed lines are cell boundaries. Scale bar is 5µm. **(B)** Graphs showing the number of *xnd-1* transcripts in P_4_ blastomeres and number of translating mRNAs comparing control and heat-shock conditions. Each vertical line represents a P_4_ blastomere. Percents indicating the percent of mRNAs that are translating. **(C)** Graphs showing the number of mRNAs (total and translating) associated with granules or cytoplasm in individual P blastomere (P_3_ for *nos-2*, and P_4_ for *xnd-1* and *zif-1*). Each horizontal gray line represents an individual blastomere. Note the distinct X-axis scales used to accommodate the higher abundance of *nos-2* transcripts (in the larger P_3_ blastomeres) compared to *xnd-1* and *zif-1* (in the smaller P_4_ blastomeres). These data were used to create the histograms shown in Fig. 3B.

**Figure EV4:**
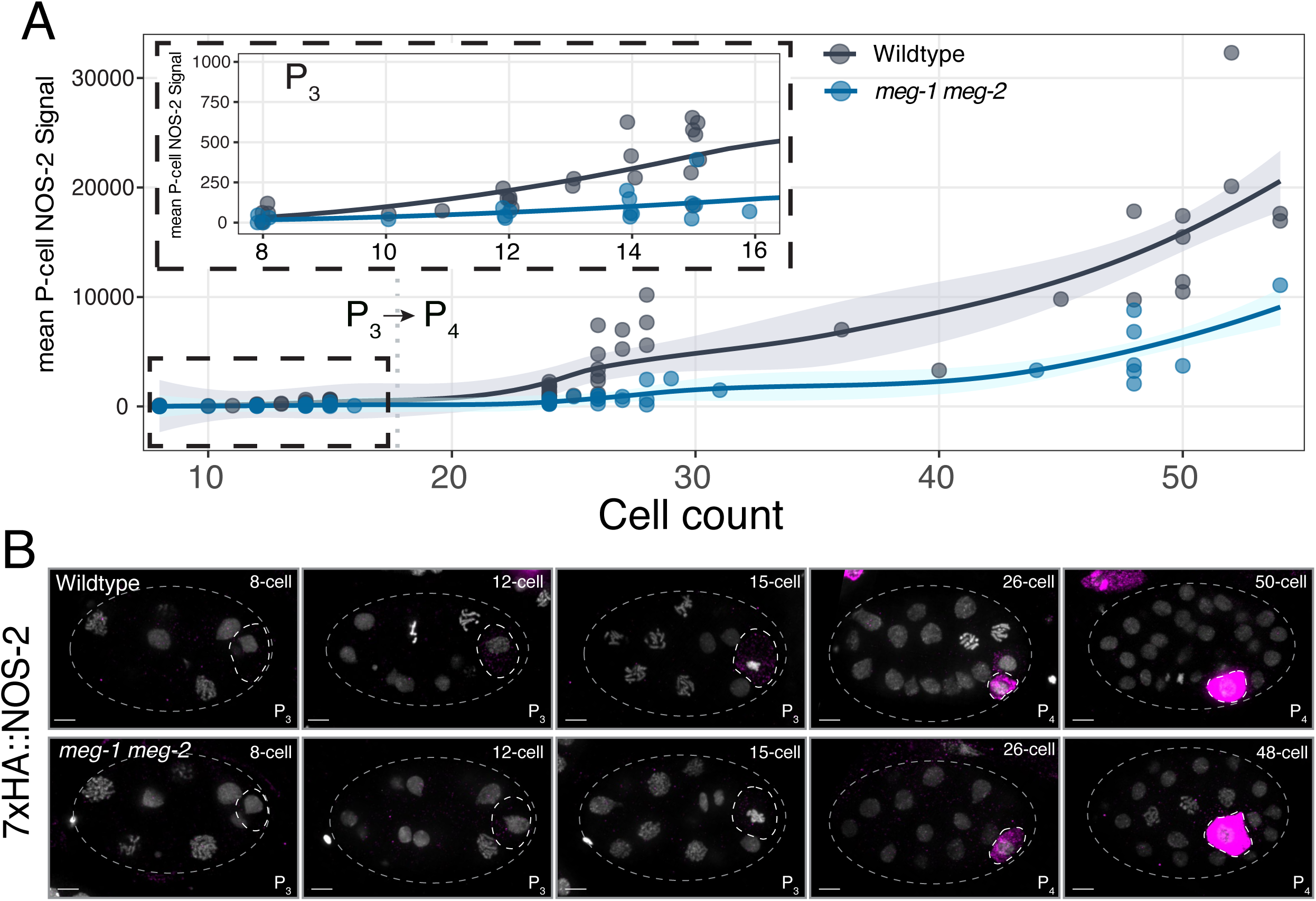
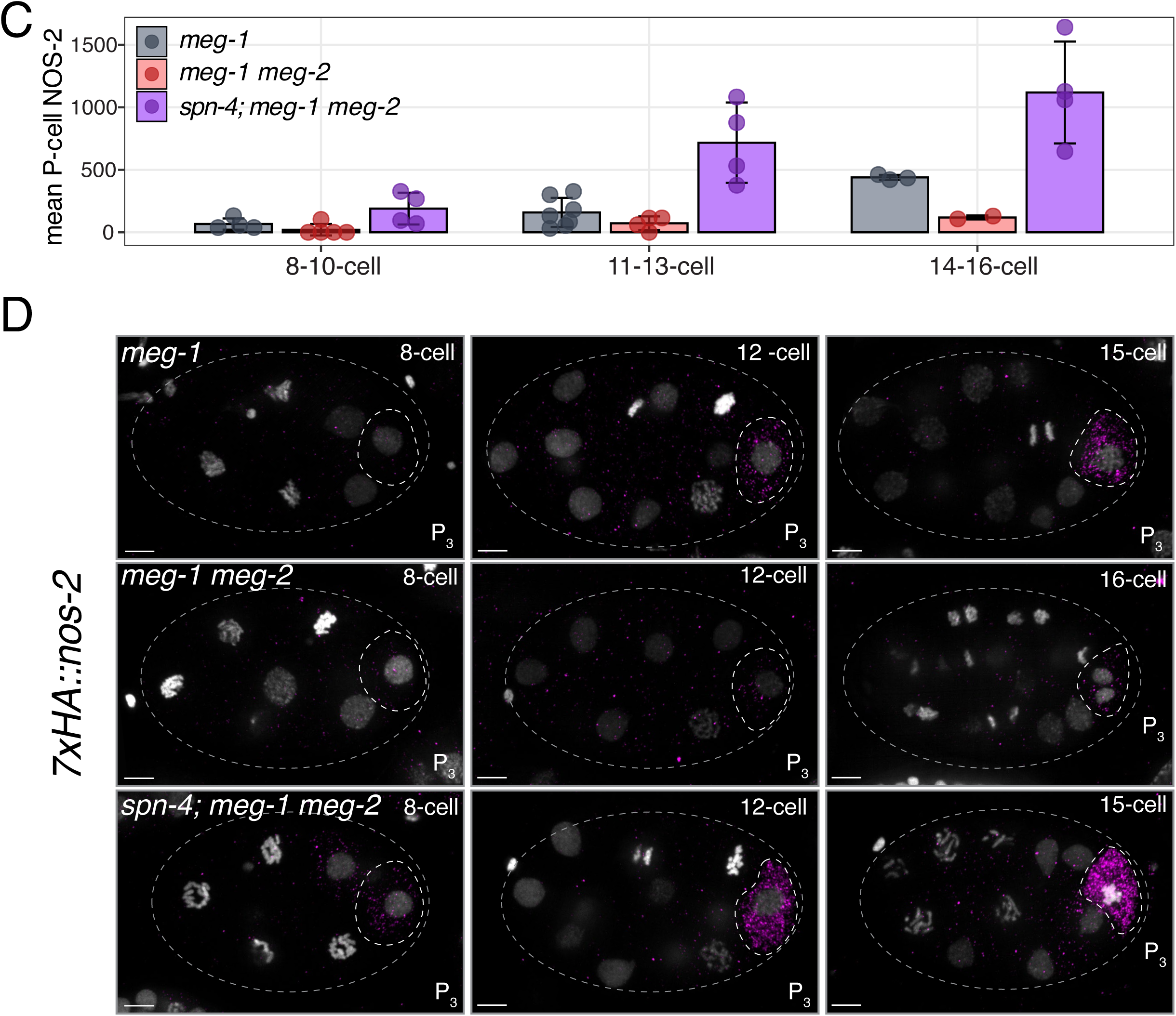
**(A)** Graph showing HA::NOS-2 signal intensity (arbitrary units, Y axis) in P blastomeres normalized to background signal in somatic blastomeres (0) comparing wild-type and *meg-1(vr10) meg-2(RNAi)* mutants at different stages (X-axis). The inset expands early stages. Each point is a single P blastomere. Lines are loess smoothed curve of the mean. Shaded areas represent 95% confidence interval. Wild-type values shown here are same as in Fig. 1B. **(B)** Photomicrographs of embryos of the indicated genotypes immunostained for 7xHA::NOS-2. **(C)** Graph showing mean 7xHA::NOS-2 signal intensity in P_3_ blastomeres comparing embryos of the indicated genotypes. Error bars represent ±1 standard deviation from the mean. Each point is one embryo. **(D)** Representative photomicrographs of the stages quantified in C. Grey dashed lines outline each embryo and the white dashed lines outline each P blastomere. Scale bars are 5µm.

**Figure EV5:**
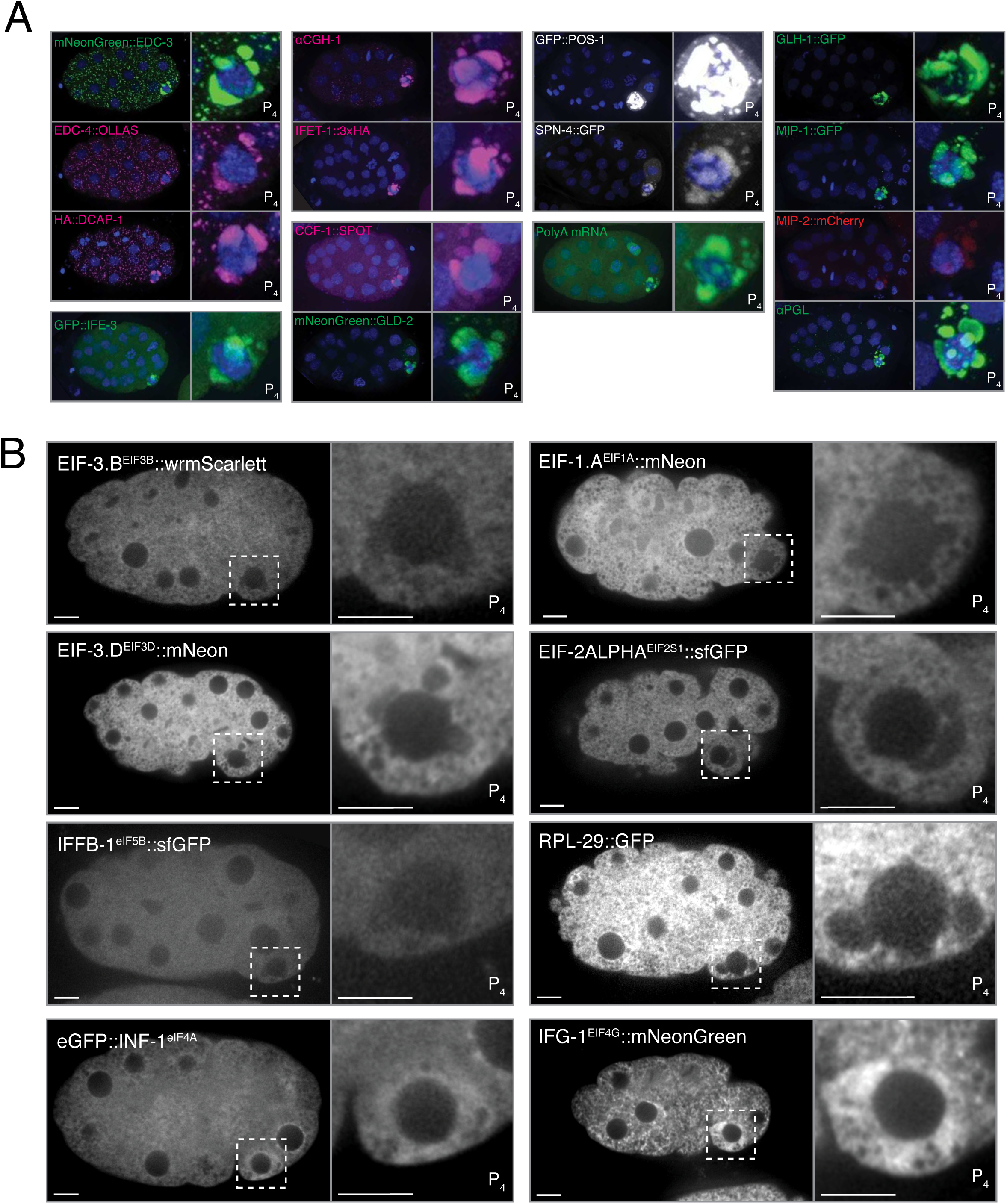
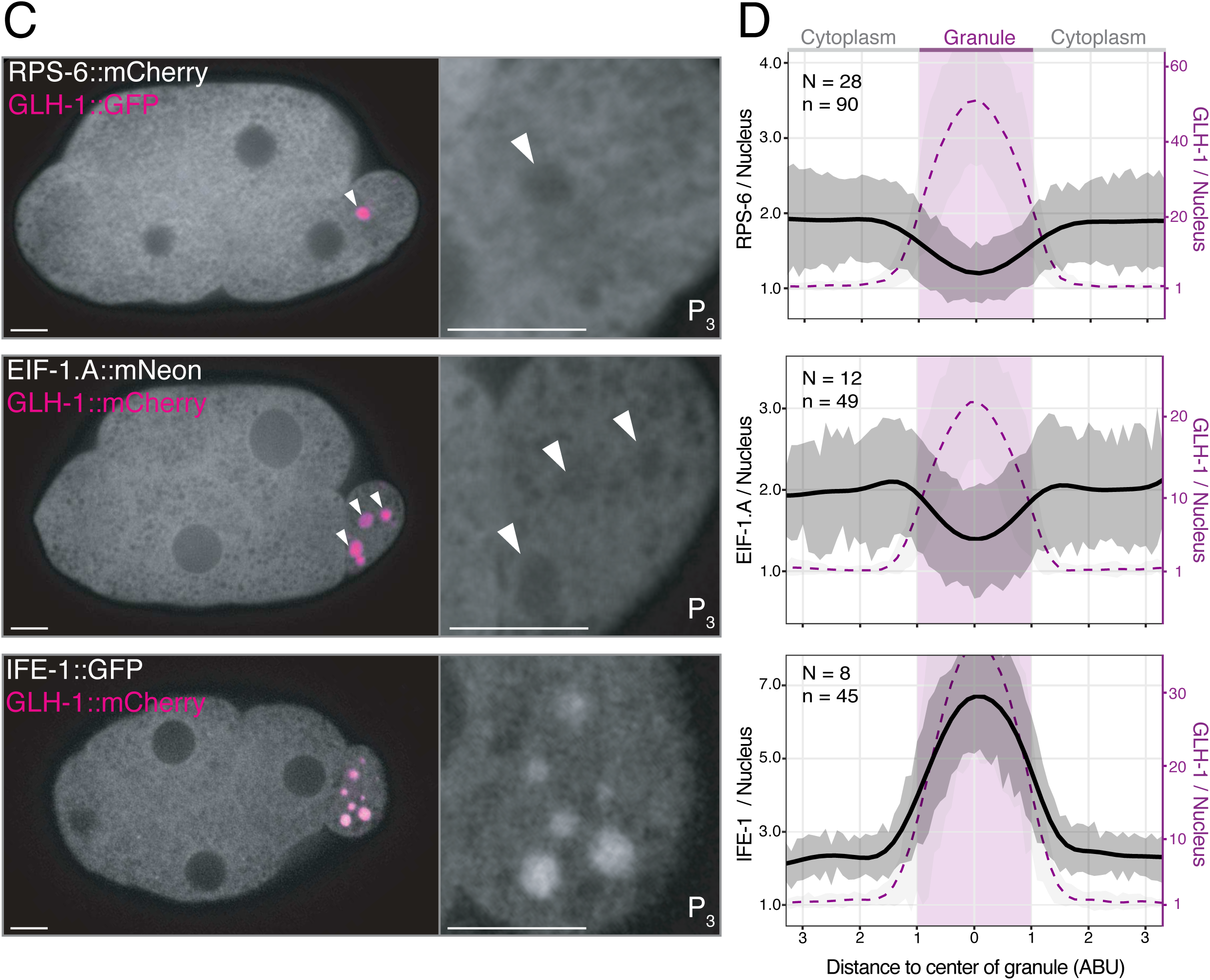
**(A)** Photomicrographs showing maximum projections of wildtype 24-to-60 cell stage embryos expressing the indicated tagged proteins visualized live (EDC-3, IFE-3, GLD-2, POS-1, SPN-4, GLH-1, MIP-1, and MIP-2) or immunostained with anti-OLLAS, anti-SpotTag, ant-HA, anti-CGH-1, or KT3 antibodies. Poly(A) smFISH shows bulk mRNA enrichment in germ granules. **(B)** Photomicrographs of single planes of live 24-60 cell stage embryos expressing the indicated fluorescent proteins. White dashed boxes mark the P_4_ blastomere, which is also shown at higher magnification. Tagged fusion proteins were expressed from their endogenous locus except for RPL-29::GFP, which was expressed from a transgene. Scale bars are 5µm. **(C)** Photomicrographs of live embryos expressing a tagged GLH-1 (GFP or mCherry, magenta) and RPS-6::mCherry, EIF-1.A::mNeonGreen, or IFE-1::GFP (gray). Arrowheads indicate P granules. Scale bars represent 5µm. Insets show a single plane photomicrographs of the P blastomere from the embryo shown without the GLH-1 channel. Scale bars are 5µm. **(D)** Graphs showing average fluorescence intensities of line scans centered on cytoplasmic P granules in P_2_ and P_3_ blastomeres for the fusion proteins shown in C. X-axis values were normalized to P granule radius based on full width at half maximum (Methods). Left Y-axis represents the enrichment of the indicated protein relative to average nuclear signal (y = 1). Right Y axis represents GLH-1 enrichment relative to average nuclear signal. Black curve represents a generalized additive model (GAM) of mean signal of target proteins along the line scans, and the dark shading surrounding it represents +/- 1 standard deviation mean. Magenta trace represents a GAM model mean of GLH-1 signal. The light gray shading around this curve represents +/- 1 standard deviation from the mean. GLH-1 enrichment values differ from those in Figure 5 due to the use of a different normalization method (Materials and Methods).

**Fig. S1:**
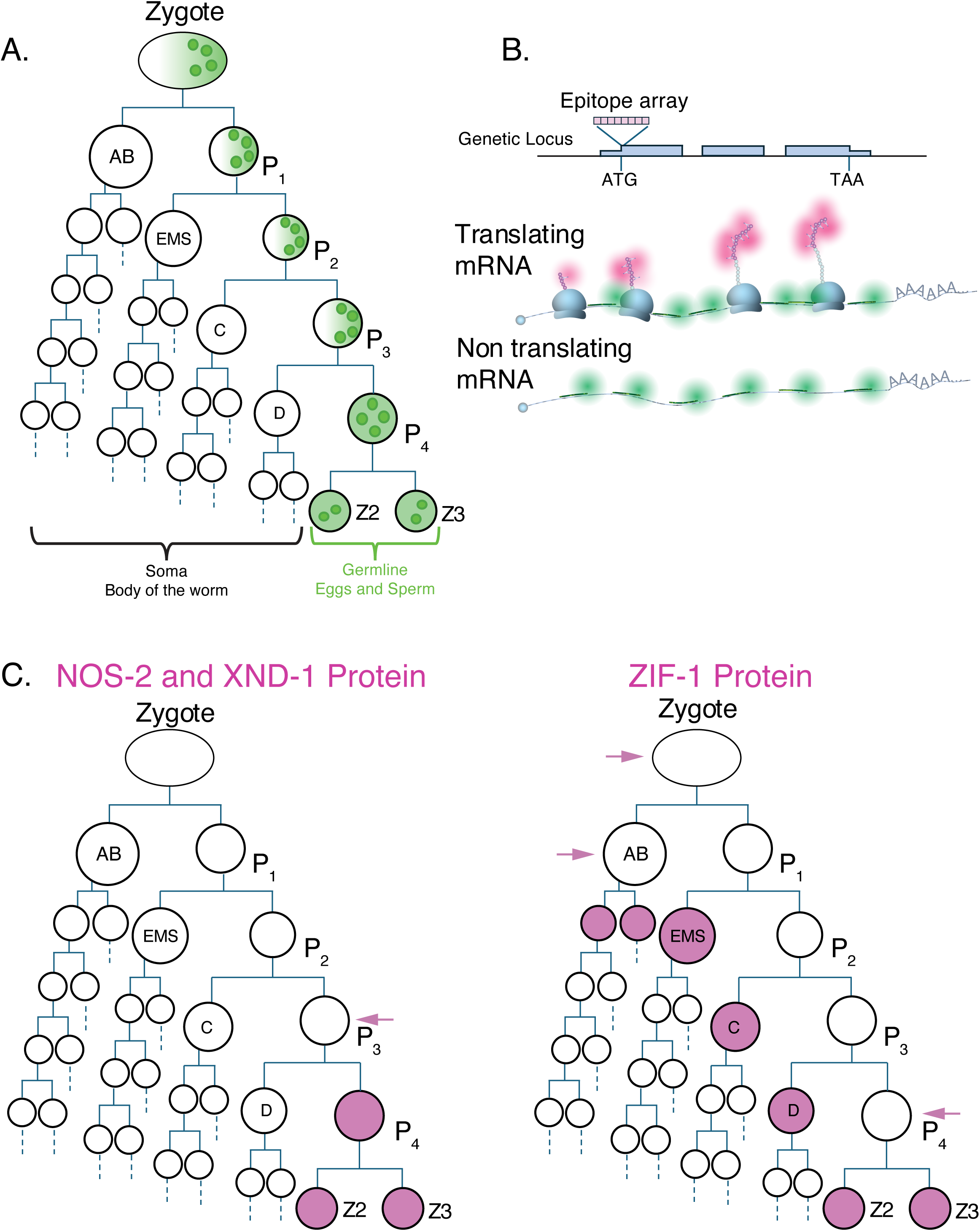
*C. elegans* lineage, tagging strategy, and expression patterns of NOS-2, XND-1 and ZIF-1. **(A)** Schematic showing the first ∼3 hours of the *C. elegans* embryonic lineage. Circles indicate cells and horizontal lines indicate cell divisions (not all divisions are shown). The zygote and P_1_-P_3_ divide asymmetrically to give rise to somatic (AB, EMS, C and D) and germline (P_1_-P_4_) blastomeres. P granules and proteins distributed between the granules and a diffuse cytoplasmic pool (e.g. POS-1) enrich asymmetrically before cell division, resulting in their preferential segregation into the germline blastomeres. P_4_ divides symmetrically into the two primordial germ cells Z2 and Z3. P granules start out distributed in the cytoplasm and become progressively more perinuclear within each P blastomeres (not shown). **(B)** Idealized schematic of tagging approach. An epitope array was inserted at the genomic locus of *nos-2, xnd-1 and zif-1* using CRISPR genome engineering (Materials and Methods). The resulting transcripts code for proteins that contain the tag array at their amino-terminus. Co-localization of RNA (detected by smFISH) with protein (detected by immunostaining) identifies translating mRNAs. (**C)** Lineage schematics as in A showing the distribution of NOS-2, XND-1 and ZIF-1 proteins as reported in the literature (pink). The arrows indicate earlier stages where translation was detected (Fig. 1 and EV1) using the tagging strategy described in B.

